# AR-V7 Utilizes a Noncanonical Nuclear Localization Signal to Sustain Androgen-Independent Nuclear Import and Signaling

**DOI:** 10.64898/2026.09.23.753944

**Authors:** Urko del Castillo, Naira E. Abou-Ghali, Colin Burdette, Michelle Naidoo, CheukMan C Au, Kiran Kumari Sahu, Xuanrong Chen, Xi Kathy Zhou, Jacob Geri, Paraskevi Giannakakou

## Abstract

AR-V7 is the most prevalent androgen receptor splice variant in metastatic castration-resistant prostate cancer, yet how it enters the nucleus independently of androgen binding has remained unclear. We previously demonstrated that AR-V7 undergoes nuclear import via a non-canonical pathway distinct from the microtubule-based, importin-α/β- and Ran-dependent mechanism used by full-length AR. Here, using systematic truncation and alanine-scanning mutagenesis, we identify a non-classical nuclear localization motif at the junction of the DNA-binding domain (DBD) and cryptic exon 3 (CE3) that is required for efficient AR-V7 nuclear import. Specific basic residues within the DBD and CE3 are required not only for nuclear entry but also for transcriptional activity. Unexpectedly, mutants retaining partial nuclear localization were transcriptionally silent, suggesting that CE3 contributes to transcriptional engagement through mechanisms separate from nuclear transport. Genome-wide transcriptomic analysis of a strongly import-deficient mutant confirmed that disruption of nuclear entry abolishes the AR-V7 transcriptional program. To map compartment-specific protein interactions, we applied antibody-guided photocatalytic proximity labeling. Nuclear AR-V7 associated most prominently with both canonical BAF and PBAF SWI/SNF chromatin remodeling complexes, together with known transcriptional coregulators and DNA-repair factors, whereas the cytoplasm-restricted mutant engaged different networks centered on mRNA deadenylation and decay, cytoskeletal regulation, and chaperone systems. Together, these findings define a variant-specific NLS, separate nuclear import from CE3-dependent transcriptional competence, and identify the cytoplasmic networks that become accessible when AR-V7 nuclear entry is impaired. They reveal two mechanistically distinct vulnerabilities for suppressing AR-V7 signaling: blocking nuclear translocation and disrupting CE3-dependent transcriptional engagement within the nucleus.

**Significance Statement:** Advanced prostate cancer is treated with drugs that target the hormone pocket of the androgen receptor. Tumors escape by making a shortened active receptor that lacks this pocket, leaving it undruggable. The shortened protein must still reach the nucleus to switch genes on, so nuclear entry is a step worth attacking. We find that the splicing event creating this variant also creates the tag that carries it into the nucleus, and that it enters by an alternative route the full receptor does not use. We also identify the protein partners the variant encounters inside and outside the nucleus, and the two sets differ strikingly, raising the question of what a cytoplasmic form of this protein does.

## Introduction

Androgen receptor (AR) signaling is critical for the initiation and progression of prostate cancer (1, 2). In early-stage hormone-sensitive disease, tumor growth can be suppressed by androgen deprivation therapy and androgen receptor pathway inhibitors (ARPi) (3–8). However, therapeutic resistance inevitably develops, leading to metastatic castration-resistant prostate cancer (mCRPC), in which AR signaling frequently persists despite castrate androgen levels and AR-directed therapy (4, 9). Persistent AR activity can arise through AR gene rearrangements, amplifications, mutations, or splice variants (10–16). Among these, AR splice variants (AR-Vs) are particularly challenging because they typically lack the ligand-binding domain (LBD) yet retain constitutive transcriptional activity in the absence of androgen (12, 17).

AR splice variant 7 (AR-V7) is the most commonly detected AR-V in mCRPC tumors, and its expression is clinically associated with poor overall survival and resistance to ARPi and taxane chemotherapy (18–21). Preclinical studies have shown that targeting AR-V7 expression through spliceosome inhibition (22), AR-FL inactivation (13), or direct degradation (23, 24) can inhibit tumor growth and sensitize CRPC models to ARPi. Yet despite this proof-of-concept, attempts to specifically inhibit AR-V7 in the clinic have not succeeded. Because AR-V7 lacks the LBD, it is intrinsically resistant to AR antagonists that bind this domain as well as to androgen-synthesis inhibitors that act by depleting its ligand. Moreover, its large intrinsically disordered N-terminal domain and DNA-binding domain lack the well-defined ligand-binding pocket that has enabled successful pharmacological targeting of AR-FL, leaving AR-V7 effectively undruggable by conventional approaches. One of the few tractable points of intervention for such a variant is its nuclear import. Because AR-V7 must enter the nucleus to drive transcription, preventing its nuclear entry offers a strategy to neutralize it without directly targeting the receptor itself.

Inhibition of AR-FL nuclear translocation has received considerable attention as a therapeutic strategy (3) and has been exploited clinically with taxanes. For AR-FL, the mechanisms governing nuclear import are well characterized (25–27). Hormone binding and Hsp90-dependent chaperone cycling (28) expose a conserved bipartite nuclear localization signal (NLS), which is recognized by importin-α/β (26, 29) and mediates passage through the nuclear pore complex in a Ran-GTP-dependent manner (25). We previously showed that AR-FL associates with microtubules, which facilitate its nuclear trafficking (30) and that taxanes can inhibit AR nuclear accumulation by disrupting this process. Importantly, we subsequently established the clinical relevance of this mechanism in the TAXYNERGY trial, demonstrating in patient samples that taxane treatment impaired AR nuclear accumulation and that this effect was associated with clinical response (31).

AR-V7, however, presents a fundamentally different trafficking scenario. Alternative splicing removes the hinge region, which contributes to the bipartite NLS and mediates microtubule binding (32), while introducing a unique 16 amino-acid C-terminal sequence encoded by cryptic exon 3 (CE3) (12, 17). The net result is a variant that localizes to the nucleus constitutively, independently of androgen binding (12), microtubules (32), or importin-α/β (33).

Previous mutagenesis work implicated basic residues in the DBD (R617/K618) and CE3 domains (K629/R631) in AR-V7 nuclear localization, as alanine substitutions at these positions caused partial cytoplasmic retention and reduced transcriptional activity (28). However, the complete architecture of the AR-V7 NLS, the mechanism by which it bypasses the canonical import machinery, and whether CE3 contributes to AR-V7 function beyond facilitating nuclear entry all remained undefined. Moreover, because AR-V7 has been studied predominantly as a nuclear transcription factor (34), little is known about the protein networks it encounters in the cytoplasm during its transit to the nucleus. Because protein interaction networks are themselves becoming druggable, knowing which partners AR-V7 engages, and in which compartment, has direct therapeutic relevance.

Here, we sought to define the molecular determinants of AR-V7 nuclear import and determine how its subcellular localization shapes its functional protein environment. Combining systematic truncation and alanine-scanning mutagenesis with genome-wide transcriptomics and antibody-guided photocatalytic proximity labeling, we map a non-classical, largely Ran-independent nuclear localization signal to the DBD-CE3 junction, uncover a requirement for CE3 residues in transcriptional activity beyond nuclear import, and define distinct protein proximity networks associated with nuclear and cytoplasmic AR-V7. Together, these findings reframe AR-V7 as a multifunctional protein whose biology extends beyond nuclear transcription and identify nuclear import and CE3-dependent transcriptional activity as distinct potential therapeutic vulnerabilities.

## Results

### AR-V7 nuclear translocation is largely independently of Ran and is conferred by the DBD-CE3 region

Our previous work demonstrated that AR-V7 employs a nuclear import mechanism distinct from that of AR-FL (33). Unlike AR-FL, which requires ligand binding and importin-αβ for nuclear translocation, AR-V7 accumulates in the nucleus constitutively through an incompletely characterized pathway. To dissect the differences in these pathways, we expressed EGFP-tagged AR-FL or AR-V7 under doxycycline-inducible control in HEK293T cells, selected for their minimal endogenous AR expression and high transfection efficiency. Cells were maintained in charcoal/dextran-stripped serum to reduce endogenous steroids, enabling precise control of ligand-dependent responses by adding the synthetic androgen R1881. We first confirmed that this cell system recapitulates the established nuclear import properties of both AR-FL and AR-V7 (SI Appendix, Fig. S1). Fixed-cell confocal microscopy confirmed the expected localization of both receptor proteins. EGFP-AR-FL was predominantly cytoplasmic without ligand and translocated to the nucleus upon R1881 treatment, whereas EGFP-AR-V7 remained nuclear even in the absence of hormone (SI Appendix, Fig. S1B and C). To determine the dependence of AR-FL and AR-V7 nuclear trafficking on the Ran GTPase cycle, we co-expressed dominant-negative GTP hydrolysis-deficient mCherry-RanQ69L into these cells (35). Expression of RanQ69L significantly impaired ligand-induced EGFP-AR-FL nuclear accumulation, from 66% to 31% nuclear signal, while modestly reducing EGFP-AR-V7 nuclear accumulation, from 70% to 60% (SI Appendix, Fig. S1D and E). AR-V7 therefore enters the nucleus largely independently of the Ran cycle that controls AR-FL localization. This is consistent with the conclusion that AR-V7 nuclear import pathway is distinct from classical Ran-dependent nuclear transport.

To define the molecular basis of this alternative import mechanism, we mapped nuclear localization determinants within AR-V7. Structurally, AR-V7 retains the N-terminal domain (NTD) and DNA-binding domain (DBD) of AR-FL but terminates immediately before the hinge domain, and instead acquires a unique 16-amino-acid sequence encoded by cryptic exon 3 (CE3) (SI Appendix, Fig. S1A). Critically, AR-V7 lacks the hinge region that contains part of the bipartite NLS required for AR-FL nuclear import, suggesting that AR-V7 relies on a distinct nuclear localization signal. We first assessed whether individual AR-V7 domains possess autonomous nuclear-localizing activity. All constructs were EGFP-tagged to enable quantitative analysis of nuclear-cytoplasmic distribution by confocal microscopy. EGFP-AR-V7 served as positive control, showing strong nuclear accumulation (87 ± 8% nuclear signal; Fig. 1B and C). EGFP alone can passively equilibrate across nuclear pores because of its small size (27 kDa) (36, 37), showed 49 ± 10% nuclear signal and therefore provided a baseline for a protein without net nuclear enrichment (38). For this mapping analysis, we operationally defined an import-competent construct as one whose mean nuclear localization fell within 1.96 standard deviations of the AR-V7 mean measured in the same experiment, corresponding to a threshold of 71.4% for the constructs in Fig. 1C. EGFP-tagged constructs containing the isolated NTD, DBD, or CE3 domains each remained predominantly cytoplasmic in HEK293T cells, with mean nuclear signals of 41%, 51% and 55%, respectively, similar to the 49% EGFP baseline and well below the threshold (Fig. 1B and C). These results indicated that no single AR-V7 domain is sufficient for nuclear targeting. We therefore generated NTD-DBD and DBD-CE3 truncation mutants. The DBD-CE3 fragment (amino acids 556-643) displayed strong nuclear localization (84 ± 10% nuclear signal), approaching that of full-length AR-V7 and well above the import-competence threshold, whereas NTD-DBD remained predominantly cytoplasmic (mean nuclear signal of < 50%) (Fig. 1B and C). These findings identify the DBD-CE3 fragment as the minimal sequence sufficient for AR-V7 nuclear import. To ask whether import competence was accompanied by transcriptional activity, we measured androgen response element (ARE)-driven luciferase reporter activity for AR-V7 and the domain constructs (Fig. 1E). AR-V7 drove strong reporter activity, whereas none of the domain constructs was transcriptionally active. Notably, the DBD-CE3 fragment was inactive despite recapitulating AR-V7 nuclear import, consistent with the absence of the N-terminal AR transactivation domain. Together, these data identify DBD-CE3 as the region where the unique signal may be located that mediates the transport of AR-V7 into the nucleus.

**Figure 1.**
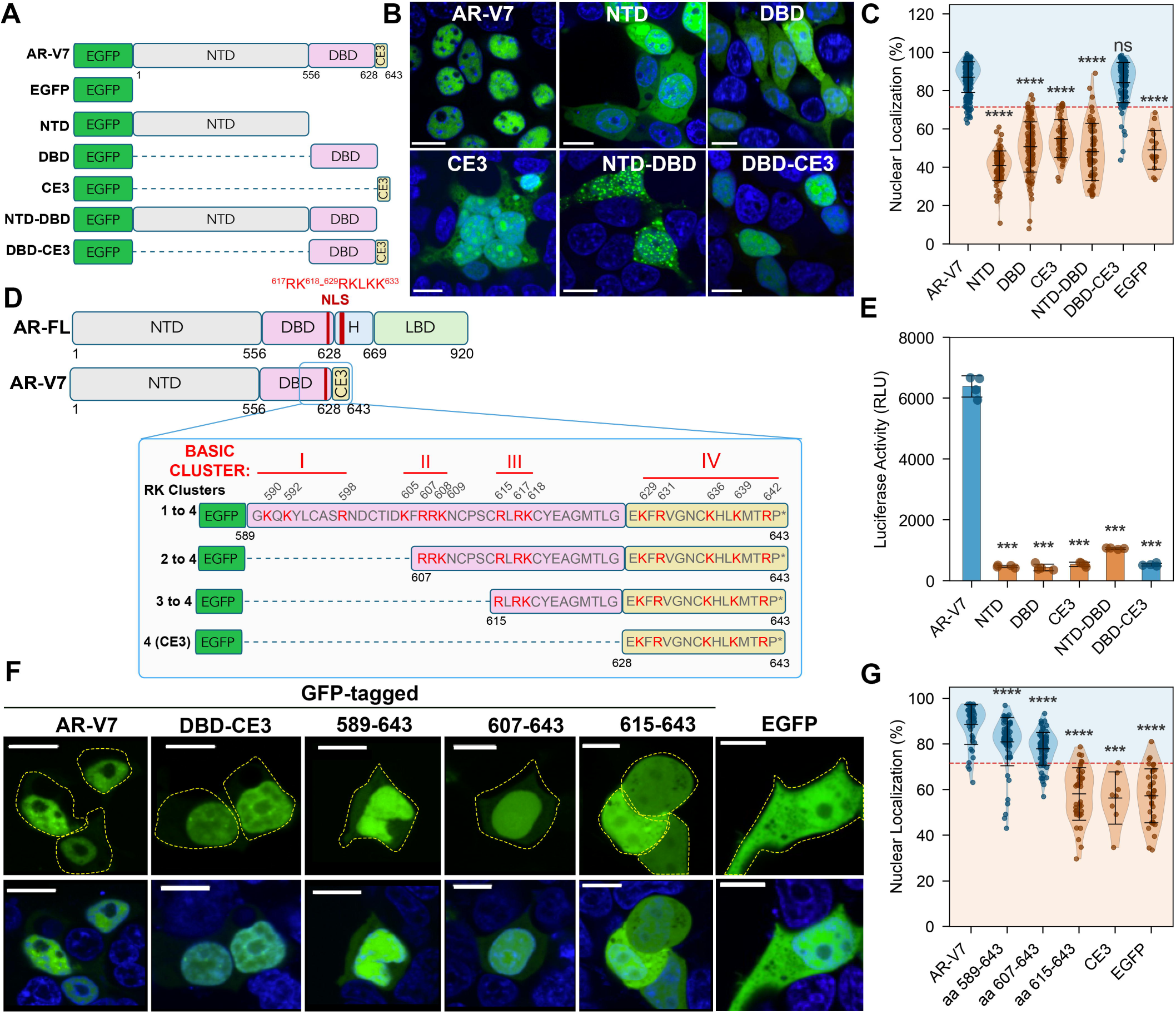
A non-classical nuclear localization signal at the DBD-CE3 junction is necessary and sufficient for AR-V7 nuclear import. **(A)** Schematic of the EGFP-tagged AR-V7 constructs used to map autonomous nuclear import activity: full-length AR-V7, EGFP alone, the isolated N-terminal domain (NTD), DNA-binding domain (DBD), and cryptic exon 3 (CE3), and the NTD-DBD and DBD-CE3 fragments. **(B)** Representative confocal images of HEK293T cells expressing the indicated EGFP-tagged domain constructs. Green, EGFP; blue, Hoechst. Scale bars, 10 µm. **(C)** Percentage nuclear localization (nuclear/whole-cell integrated EGFP intensity) for each domain construct. Data points are individual cells overlaid on violin plots; lines indicate mean ± SD. The dashed red line marks the import-competence threshold (AR-V7 mean-1.96xSD, 71.4%); constructs whose mean nuclear localization falls above the line, in the green-shaded region, are scored import-competent, whereas those falling below it, in the orange-shaded region, are scored import-defective. Each construct was compared to AR-V7 by two-sided Mann-Whitney U test with Bonferroni correction for six comparisons. Only the DBD-CE3 fragment recapitulated AR-V7 nuclear accumulation. **(D)** Top, domain architecture of AR-FL and AR-V7 highlighting the DBD-CE3 region that contains the NLS. Bottom, sequence of the DBD-CE3 junction (aa 589-643) showing four clusters of basic residues (I-IV; basic residues in red) and the EGFP-tagged peptides used for fine-mapping: Clusters I-IV (aa 589-643), Clusters II-IV (aa 607-643), Clusters III-IV (aa 615-643), and Cluster IV alone, corresponding to CE3 (aa 628-643). **(E)** Androgen-response-element (ARE)-driven luciferase reporter activity (RLU) for AR-V7 and the domain constructs. Bars indicate the mean of four independent experiments, each performed in technical triplicate; points are the individual experiment means. Each construct was compared to AR-V7 by unpaired Welch’s t-test on the four experiment means, with Bonferroni correction for five comparisons. **(F)** Representative confocal images of HEK293T cells expressing EGFP alone, EGFP-AR-V7, EGFP-DBD-CE3, or the EGFP-tagged fine-mapping peptides (aa 589-643, 607-643, 615-643). Top, EGFP; bottom, merge with Hoechst. Dashed outlines indicate the cell periphery. Scale bars, 10 µm. **(G)** Percentage nuclear localization for the truncation peptides, pooled from two independent experiments, with shading as in (C). The import-competence threshold (AR-V7 mean - 1.96 × SD) is 71.6%. Significance versus AR-V7 was determined using simultaneous tests of general linear hypotheses (Bonferroni-adjusted for five comparisons) applied to a linear mixed-effects model with construct as a fixed effect, experiment as a random intercept, and construct-specific variances; n indicates individual cells. The minimal import-competent peptide was aa 607-643. *p < 0.05; **p < 0.01; ***p < 0.001; ****p < 0.0001; ns, not significant.

### Basic residues in Cluster II-IV of the DBD-CE3 junction define a non-classical NLS in AR-V7

To identify the specific sequence determinants mediating nuclear import, we examined the distribution of basic residues within the DBD-CE3 region. Basic amino acids (lysine and arginine) are hallmark features of nuclear localization signals (39). Whereas classical NLS motifs engage importin-α/β through Ran-dependent mechanisms, basic residue clusters can also drive carrier- or RanGTP-independent import by forming electrostatic surfaces that contact FG-repeat nucleoporins directly, as demonstrated for transcription factors (40, 41), ribosomal proteins (42), and viral oncoproteins (43). Sequence analysis revealed four distinct basic residue clusters spanning amino acids 589-643 at the DBD-CE3 junction (Fig. 1D). These clusters lie within and adjacent to the second zinc finger domain (ZFD) and encompass the dimerization box motif (YLCASRNDCTI), which we previously demonstrated to be involved in AR-V7 nuclear localization through mutations at positions A596 and S597 (33). Their consecutive alternating arrangement suggested they may form a non-canonical NLS, as previously identified for other transcription factors (39, 44, 45). To determine which basic residue clusters are required for nuclear import, we generated a panel of EGFP-tagged peptides encoding different combinations from within the DBD-CE3 domain. These comprised of Clusters I-IV (aa 589-643), Clusters II-IV (aa 607-643), Clusters III-IV (aa 615-643), and Cluster IV alone (aa 628-643), which corresponds to the CE3 domain (Fig. 1D). Live-cell imaging revealed a clear hierarchy of import efficiency. Peptides spanning residues 589-643 and 607-643 accumulated in the nucleus above the import-competence threshold of 71.6% (81% and 78%, respectively), as did full-length AR-V7 (89%) and the complete DBD-CE3 fragment (Fig. 1F and G). In contrast, the truncated peptide 615-643 fell below the import-competence threshold (58%), as did CE3 alone (56%). Neither was distinguishable from free EGFP (57%), indicating no nuclear enrichment above free diffusion (Fig. 1G). These results establish residues 607-643 as the minimal NLS of AR-V7, with upstream sequences (589–606) potentially enhancing import efficiency.

### Alanine scanning defines the AR-V7 NLS and reveals CE3-dependent transcriptional requirements

To determine the exact residues required for nuclear import, and to test whether they could be disrupted without compromising the rest of the AR-V7 protein, we performed alanine-scanning mutagenesis across basic Clusters II-IV in full-length AR-V7 (Fig. 2A). These constructs are full-length AR-V7 carrying point substitutions, so the N-terminal transactivation domain remains intact and any loss of reporter activity cannot be explained by removal of the transactivation machinery. Constructs are named Dx/Cy, where x and y are the numbers of substitutions in the DBD and in CE3, respectively. To bound the assay, we normalized nuclear localization to two references measured by quantitative confocal microscopy. Constitutively nuclear AR-V7 (87.3% of total cellular AR in the nucleus) and cytoplasmic unliganded AR-FL (56.6%) (SI Appendix, Fig. S2) were set to 100% and 0%, respectively. All mutants are reported on this scale as linear mixed-effects estimates. Ligand-treated AR-FL, a positive control for import, recovered to 74% of AR-V7 (Fig. 2B and D). Previous work showed that substitutions in the DBD R617A/K618A contributes to nuclear localization, and that the D2/C2 mutant, which adds the CE3 substitutions K629A/R631A, partially reduces AR-V7 nuclear accumulation (28). We confirmed this partial phenotype, with D2/C2 retaining a raw nuclear fraction of 65%, corresponding to 29% of AR-V7, a ∼70% loss of nuclear localization (Fig. 2B and E). Using D2/C2 as a backbone, we progressively extended the substitutions to K636A (D2/C3), K636A/K639A (D2/C4), and K636A/K639A/R642A (D2/C5) (Fig. 2A). Each addition further compromised localization, to 14% (D2/C3), 10% (D2/C4), and 2% (D2/C5), all significantly below AR-V7, confirming a role of CE3 in AR-V7 nuclear translocation (Fig. 2B and E).

**Figure 2.**
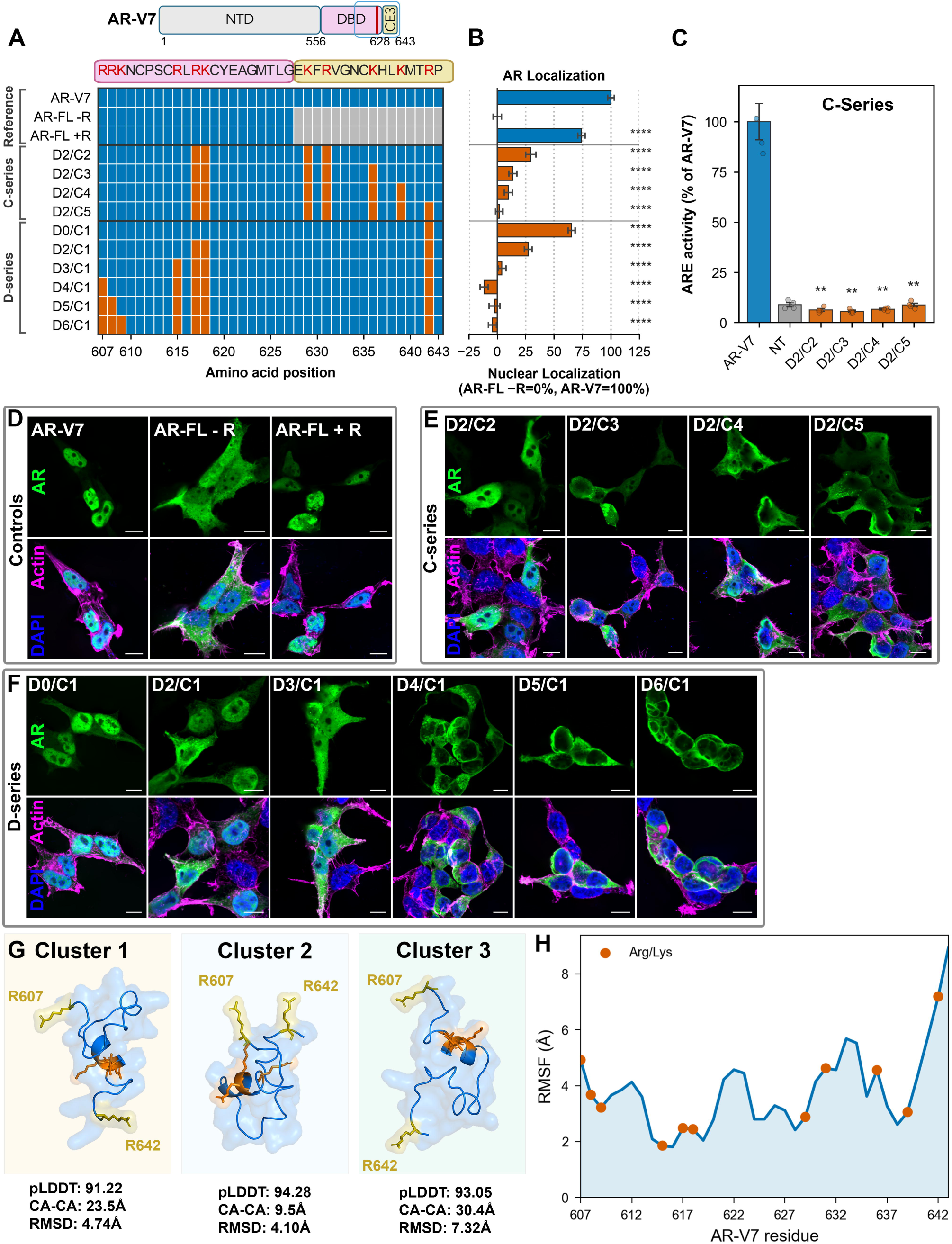
Alanine-scanning mutagenesis defines the core NLS residues and reveals a conformationally flexible signal. **(A)** Map of the alanine-scanning mutant panel. Top, sequence of the DBD-CE3 junction (aa 607-643) with basic residues (Arg/Lys) in red. Each row is a construct; orange cells indicate residues mutated to alanine. Rows are grouped as Controls (AR-V7, AR-FL -R, AR-FL +R), the CE3-extending C-series (D2/C2, D2/C3, D2/C4, D2/C5), and the DBD-extending D-series (D0/C1, D2/C1, D3/C1, D4/C1, D5/C1, D6/C1). AR-FL lacks the CE3 sequence (positions shown in grey). **(B)** Percentage nuclear localization for each construct, normalized to controls (AR-FL -R = 0%, AR-V7 = 100%). Bars indicate the model-estimated mean ± SE. Significance versus AR-V7 was determined using simultaneous tests of general linear hypotheses (Bonferroni-adjusted) applied to a linear mixed-effects model with genotype as a fixed effect, experiment as a random intercept, and genotype-specific variances. AR-FL -R and AR-V7 define the 0% and 100% points of this scale. Between-mutant contrasts quoted in the Results come from the same model, Bonferroni-adjusted across the 13 contrasts tested. Raw values in Fig. S2. **(C)** ARE-driven luciferase reporter activity for the C-series mutants, expressed as a percentage of AR-V7. Bars indicate mean ± SEM; NT, non-transfected control. Significance versus AR-V7 was assessed by unpaired Welch’s t-test with Bonferroni correction for four comparisons. **(D)** Representative confocal images of the control constructs (AR-V7, AR-FL -R, AR-FL +R). Green, EGFP-tagged AR; magenta, F-actin; blue, DAPI. Scale bars, 10 µm. **(E)** Representative confocal images of the C-series mutants; channels as in (D). **(F)** Representative confocal images of the D-series mutants; channels as in (D). **(G)** Representative structures of the three conformational clusters of the AR-V7 NLS peptide (aa 607-643) from a 100-model AlphaFlow ensemble, each shown as its highest-confidence conformer. Values printed beneath each structure are those of the conformer shown. The Arg607 and Arg642 termini are highlighted**. (H)** Per-residue backbone (Cα) root-mean-square fluctuation (RMSF, Å) across the NLS peptide (residues 607-643) over the ensemble; basic residues (Arg/Lys) are marked in orange. A structured central core (residues 614-619, RMSF 1.81-2.48 Å) is flanked by flexible termini, with maximal flexibility at the C-terminus (Pro643, 8.95 Å; Arg642, 7.19 Å). *p < 0.05; **p < 0.01; ***p < 0.001; ****p < 0.0001; ns, not significant.

We next asked whether these losses track transcriptional output, using an ARE-driven Gaussia luciferase reporter (46). Wild-type AR-V7 drove strong reporter activity and was set to 100%, while non-transfected cells (NT) gave 8.9% defining assay background. All four C-series mutants were transcriptionally inactive, with values at NT (6.3%, 5.6%, 6.7% and 8.7%; Fig. 2C). Transcriptional output therefore did not track residual nuclear localization across this series, which spans 29.5% to 1.7% of AR-V7. Because the substituted residues lie within the DBD and CE3, reduced reporter activity may also reflect impaired DNA binding, which this assay cannot separate from a transcriptional role. The data therefore show that nuclear localization is necessary but not sufficient for AR-V7 transcriptional activity.

D2/C5 and D2/C4 differ only by R642A, and D2/C5 was more strongly excluded, implicating R642 in the NLS but leaving open whether it acts alone or together with upstream basic residues in Clusters II and III. To address this, we held the CE3 substitution fixed at R642A and progressively added DBD substitutions (D-series; Fig. 2A, B and F). R642A alone (D0/C1) reduced localization to 65%, and adding R617A/K618A (D2/C1) further reduced localization to 27%. Neither reached the full exclusion of D2/C5, consistent with our earlier mapping, in which 607-643 supported import whereas 615-643 did not (Fig. 1G). Adding R615A (D3/C1) drove localization down by 4%, a 23-point decrease. Both DBD additions therefore produced substantial losses on an R642A background, indicating contributions to nuclear import that are non-redundant with R642 D3/C1 phenocopied D2/C5, reaching the unliganded AR-FL baseline by a different combination of substitutions: D3/C1 mutates R615 and leaves K629, R631, K636 and K639 intact, while D2/C5 retains R615 but mutates these four CE3 residues. Full exclusion therefore does not require K636 and K639 once R615 is lost, though K636 does contribute on an R615-intact background (D2/C2, 29% *versus* D2/C3, 14%). Together, R615, R617, K618 and R642, account for most of the import defect. All four lie within residues 615-643, however, that peptide did not support import, pointing to additional determinants upstream of R615.

To test whether residues further upstream contribute, we progressively mutated R607, R608 and K609 on the D3/C1 background (Fig. 2B and F). Adding R607A (D4/C1) drove localization to - 12%, significantly below both D3/C1 and the unliganded AR-FL reference, indicating that this mutant is not merely import-deficient but is retained in the cytoplasm more effectively than AR-FL. Adding R608A (D5/C1, -2%) and K609A (D6/C1, -4%) produced no further decrease, and both mutants were indistinguishable from D4/C1. Because no construct was more cytoplasmic than D4/C1, we cannot distinguish whether R608 and K609 are dispensable or whether their contribution is masked by the assay floor. Thus, R607 extends nuclear exclusion beyond the four residues identified above, defining the AR-V7 NLS as a multipartite signal comprising R607, R615, R617, K618 and R642.

Having identified the core NLS residues, we asked whether the NLS peptide adopts a defined structure. We generated a 100-state conformational ensemble of residues 607-643 using AlphaFlow, a generative modeling approach built on AlphaFold2 (47, 48). The ensemble was overall high confidence, with 81 of 100 models exceeding the pLDDT reliability threshold of 90 (SI Appendix, Fig. S3A). Hierarchical clustering identified three conformational states (SI Appendix, Fig. S3C), each shown in Fig. 2G as a single representative conformer. Cluster 2 was the most compact and highest-confidence state (n = 40, mean pLDDT 92.48, mean Arg607-Arg642 Cα-Cα distance 21.5 Å), whereas Cluster 1 formed a heterogeneous intermediate population (n = 49, 24.6 Å), and Cluster 3 comprised a small, structurally divergent minority (n = 11, 25.8 Å). Cluster 2 was significantly more compact than both other states (Mann-Whitney p = 0.003 and 0.001), whereas Clusters 1 and 3 were indistinguishable in compactness (p = 0.24). Cluster 3 was distinguished instead by its greater deviation from the reference model, with a mean RMSD of 7.70 Å compared with 4.56 Å for Cluster 1 and 3.69 Å for Cluster 2. Per-residue flexibility analysis revealed a structured central core (residues 614-619, RMSF 1.81-2.48 Å) flanked by highly flexible termini. Flexibility increased sharply across the CE3 arm reaching 7.19 Å at Arg642 and 8.95 Å at the C-terminal Pro643 (Fig. 2H and Movie S1). Consistent with this flexibility, end-to-end distance measurements between Arg607 and Arg642 varied substantially across the ensemble (Cα-Cα 23.48 ± 4.76 Å, range 9.54-37.07 Å), with even greater variability at the side-chain level (Cζ-Cζ 25.97 ± 6.32 Å, range 8.29-45.41 Å) (SI Appendix, Fig. S3B). Thus, in the absence of interactors, the NLS samples a broad continuum from compact to extended conformations rather than adopting a single fixed geometry. This conformational flexibility, particularly at R642, may provide a structural context for the cooperative nature of the NLS, in which individual basic residues make partial contributions but combinations of mutations are required for complete disruption of nuclear import.

### Transcriptomic profiling reveals complete loss of AR-V7 transcriptional output upon nuclear import perturbation

The C-series mutants were transcriptionally inactive across a range of residual nuclear localization, leaving open whether these mutations reflected a global collapse of the AR-V7 transcriptional program or selective impairment at specific loci. To distinguish between these possibilities, we performed RNA-seq after 24 h of transgene induction in LNCaP-C4-2 cells, comparing AR-V7*^NLSmut^* (D2/C5) with wild-type AR-V7 and AR-FL in the presence and absence of R1881. AR-FL expression without androgen produced minimal transcriptional change, with 3 genes upregulated and 2 downregulated (Fig. 3A). In contrast, addition of R1881 upregulated 653 genes and downregulated 642, including the canonical AR targets FKBP5, ELL2 and PGC (Fig. 3B). Ligand-independent AR-V7 produced an even broader transcriptional response, upregulating 3,159 genes and downregulating 438 (Fig. 3C). By contrast, AR-V7*^NLSmut^* was transcriptionally silent and only AR itself was significantly upregulated, reflecting transgene expression, with no other gene meeting the significance threshold (Fig. 3D). Heatmap analysis of canonical AR target genes further showed that AR-FL+R and AR-V7 both activated the classical ARE-driven program, whereas AR-FL alone and AR-V7*^NLSmut^* did not (Fig. 3E). Gene set variation analysis (GSVA) (49) across four independent AR activity signatures confirmed that AR-V7 recapitulates ligand-driven AR-FL transcriptional activity, and that this activity is entirely abolished when nuclear entry is disrupted in AR-V7*^NLSmut^* (Fig. 3F). Together, these transcriptomic data establish that disruption of these basic residues abolishes the AR-V7 transcriptional program. Combined with the C-series reporter data, in which mutants retaining partial nuclear localization were nevertheless transcriptionally inactive (Fig. 2B and C), these findings indicate that nuclear import is necessary but not sufficient for AR-V7 transcriptional activity and suggest that the mutated CE3 residues also contribute to transcriptional competence downstream of nuclear import, such as at the level of DNA binding or cofactor recruitment.

**Figure 3.**
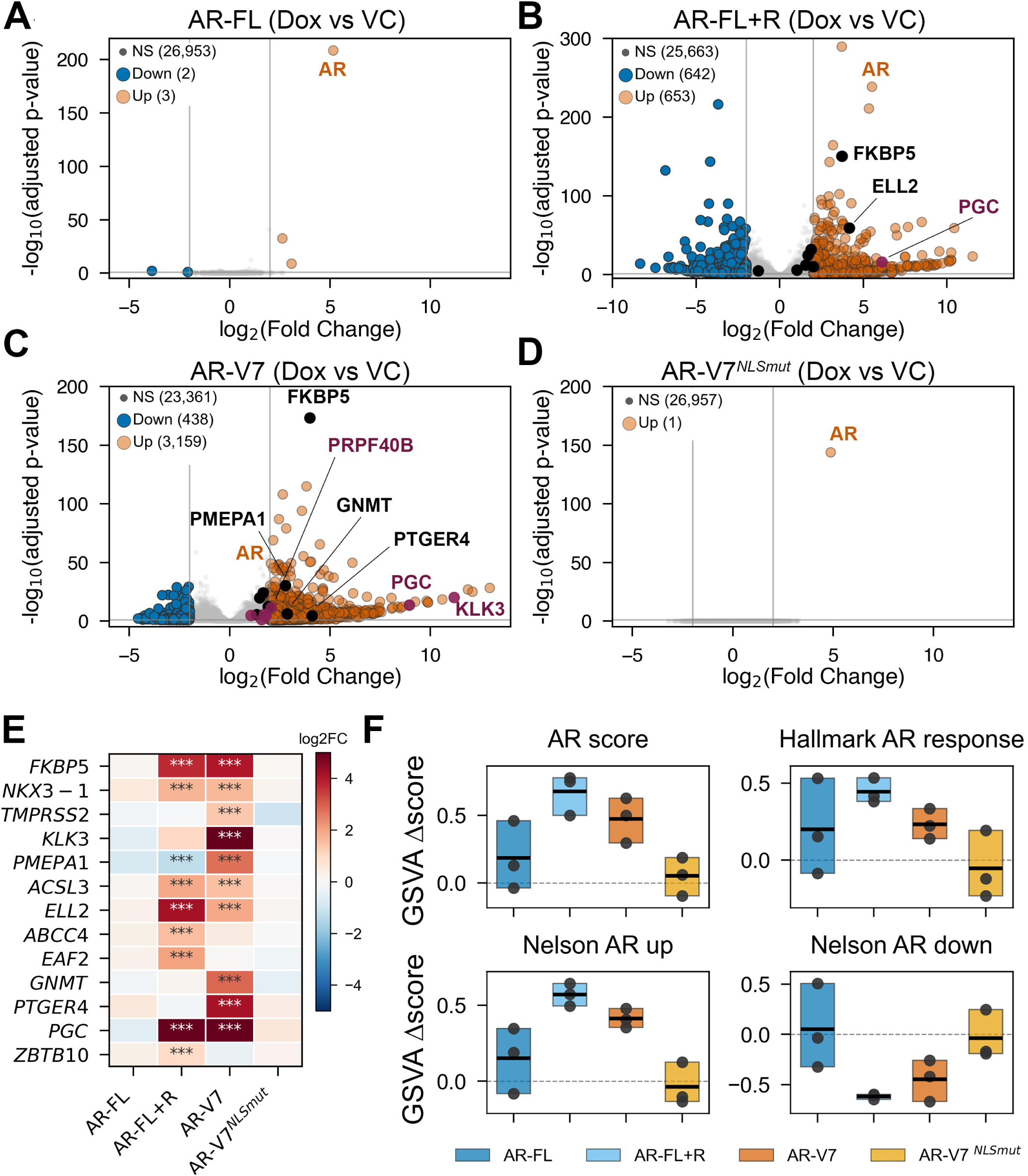
AR-V7 drives a broad androgen-responsive transcriptional program that is abolished by NLS mutation. **(A-D)** Volcano plots of differential gene expression (DESeq2) in LNCaP-C4-2 cells expressing doxycycline-inducible AR constructs (Dox) versus vehicle control (VC). Dashed lines indicate significance thresholds (|log_2_FC| > 2; Benjamini-Hochberg adjusted *p* < 0.1). Genes are classified as upregulated (orange), downregulated (blue), or not significant (grey); selected AR target genes are labeled. **(A)** AR-FL without androgen. **(B)** AR-FL with R1881, induces canonical AR targets (e.g., FKBP5, ELL2, PGC). **(C)** AR-V7 without androgen, produces the largest transcriptional footprint of the four conditions, including canonical AR targets (FKBP5, PMEPA1, GNMT, PTGER4, KLK3, PGC) and variant-associated transcripts (PRPF40B). **(D)** AR-V7^NLSmut^ (D2/C5) without androgen, transcriptionally inert except for the AR transgene itself. **(E)** Heatmap of log_2_(fold change) for canonical AR target genes across the four conditions (color scale: red, upregulated; blue, downregulated). Significance is annotated from DESeq2 adjusted *p*-values: \*\*\**p* < 0.001, \*\**p* < 0.01, \**p* < 0.05. **(F)** Gene set variation analysis (GSVA) Δscores for four independent AR activity signatures (AR score, Hallmark AR response, Nelson AR up, Nelson AR down) across AR-FL, AR-FL+R, AR-V7, and AR-V7^NLSmut^. Floating bars span the replicate range with a line at the mean; individual replicates are shown as dots (n = 3 per group). Δscore is the change in GSVA enrichment upon Dox-induced AR expression relative to the mean of the matched vehicle-control group; the dashed line indicates zero. Significance was assessed by one-sample t-test of the three per-replicate Δscores against zero within each signature and group. AR-V7 differed from zero in two of the four signatures (p = 0.008 and 0.039) and was marginal in the other two (p = 0.055 and 0.064), whereas no signature differed from zero in AR-V7^NLSmut^.

### Photocatalytic proximity labeling reveals compartment-specific AR-V7 protein networks

AR-V7 is widely studied as a constitutively active transcription factor, yet its biology in the cytoplasm remains poorly understood. Because AR-V7 is rapidly imported into the nucleus with minimal cytoplasmic residence (33), proteins that engage it before it reaches the nucleus have been largely invisible to conventional protein-protein interaction methods. The nuclear import-defective AR-V7*^NLSmut^*, which accumulates predominantly in the cytoplasm, provided a strategy to capture these otherwise transient cytoplasmic interactions by proximity labeling.

To map the protein interactome of AR-V7 and AR-V7*^NLSmut^*, we employed µMap, a carbene-based photocatalytic proximity labeling proteomics workflow (Fig. 4A-C) (50). Through the µMap platform, a target-specific primary antibody is paired with a secondary antibody conjugated to an iridium photocatalyst. Upon blue-light irradiation (440 nm), diazirine-biotin is converted into reactive carbene intermediates that covalently label proteins within approximately 4 nm of the catalyst, restricting labeling to the target and its immediate neighbors (51).

**Figure 4.**
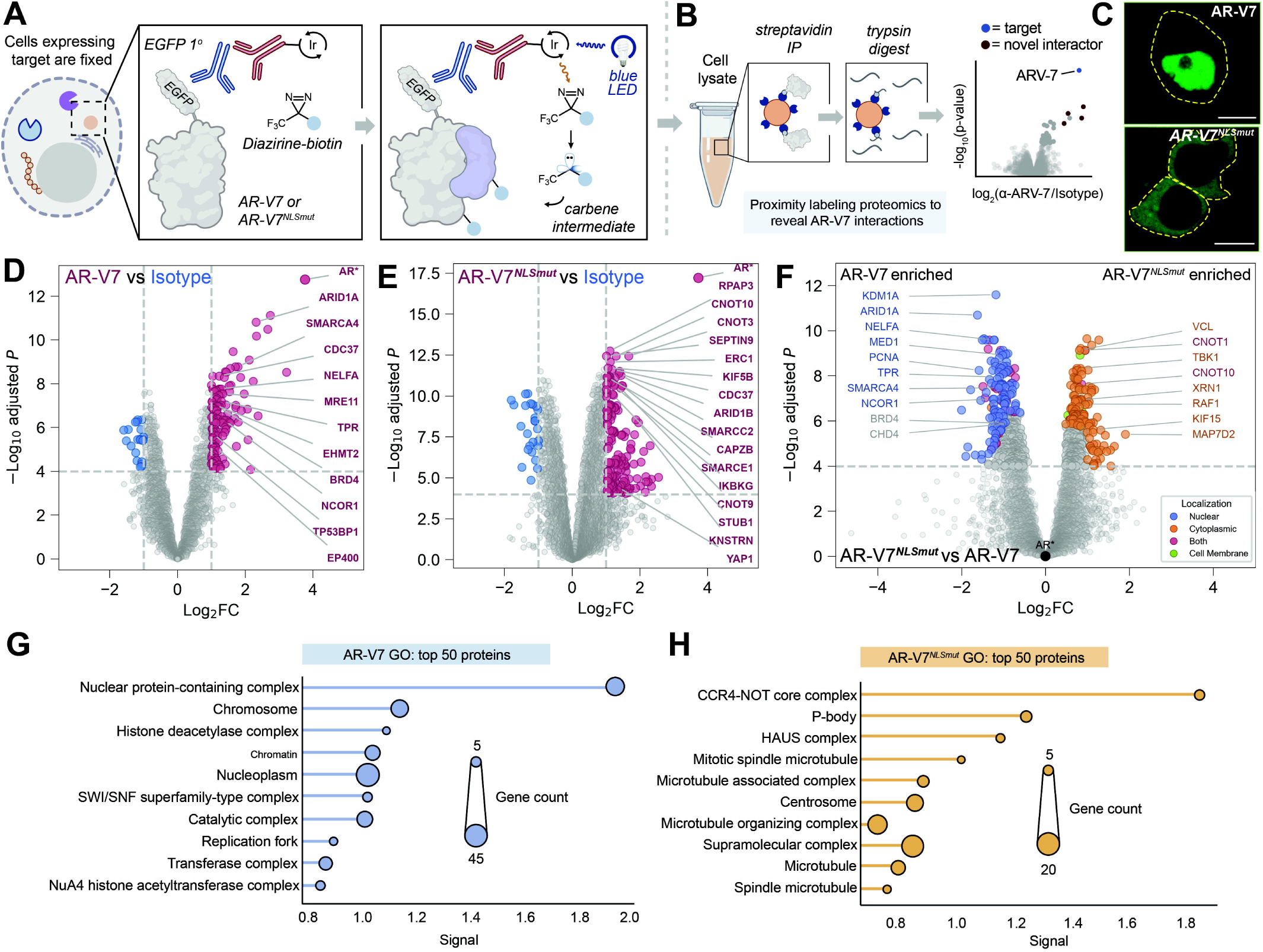
Photocatalytic proximity labeling (µMap) reveals compartment-specific AR-V7 protein proximity networks. **(A)** Schematic of µMap photocatalytic proximity labeling. In fixed cells expressing EGFP-tagged AR-V7 or AR-V7*^NLSmut^*, an anti-GFP rabbit primary antibody binds the bait; an iridium-conjugated anti-rabbit secondary antibody serves as the photocatalyst. Blue-light (440 nm) irradiation activates a biotinylated diazirine probe to a reactive carbene intermediate that covalently labels proteins within an ∼4 nm radius. **(B)** Experimental workflow: cell lysates are enriched on streptavidin beads, trypsin-digested, and analyzed by mass spectrometry; enriched proteins (bait, blue; novel interactors, red) are resolved by quantitative volcano analysis. **(C)** Confocal fluorescence microscopy of GFP-tagged AR-V7 (top) and AR-V7*^NLSmut^* (bottom) in fixed LNCaP-C4-2 cells. Yellow dashed lines indicate cell boundaries. Scale bars, 10 µm. **(D)** Volcano plot of µMap-enriched proteins for AR-V7 versus IgG isotype control (n = 10 per arm). Dashed lines indicate the significance thresholds, |log2FC| > 1 and Benjamini-Hochberg adjusted p < 1e-4. Magenta, significantly enriched; blue, significantly depleted; grey, non-significant. AR (bait) is marked with an asterisk (*). **(E)** Volcano plot of µMap-enriched proteins for AR-V7*^NLSmut^* versus IgG isotype control; thresholds and color scheme as in (D). **(F)** Differential interactome of AR-V7*^NLSmut^* versus AR-V7, from a linear model with variance-stabilizing normalization that controls for background and shared interactors. The dashed line marks the significance threshold, Benjamini-Hochberg adjusted p < 1e-4; no fold-change cutoff was applied. Significant proteins are colored by annotated subcellular localization: nuclear (blue), cytoplasmic (orange), both (red), cell membrane (green); proteins without a localization annotation are shown in grey. **(G)** Gene Ontology cellular component enrichment analysis (STRING v12) for the top 50 proteins significantly enriched in the AR-V7 interactome. Dot size indicates gene count, and the x-axis shows the STRING enrichment strength (Signal). **(H**) As in (G), for the top 50 proteins significantly enriched in the AR-V7NLSmut interactome.

We first validated µMap using a doxycycline-inducible EGFP-AR-FL model in paraformaldehyde (PFA)-fixed LNCaP-C4-2 cells. AR-FL was the most significantly enriched protein relative to IgG isotype controls (SI Appendix, Fig. S4A). Of the 218 proteins significantly enriched at log2FC > 1 and Benjamini-Hochberg adjusted p < 0.01, 22 had been reported as AR interactors in published RIME and BioID datasets (52, 53), representing a 3.6-fold enrichment over their frequency among all 4,392 proteins recovered (hypergeometric p = 1.1 x 10-7). Thus, µMap selectively recovered the AR-associated protein environment rather than abundant cellular proteins nonspecifically (SI Appendix, Fig. S4A).

We next applied µMap to EGFP-AR-V7 expressed in LNCaP-C4-2 cells. Relative to isotype controls, 129 proteins were significantly enriched at |log2FC| > 1 and Benjamini-Hochberg adjusted p < 1e-4, and this wild-type AR-V7 interactome was dominated by nuclear proteins (Fig. 4D), with enriched factors spanning chromatin remodeling, transcription elongation, epigenetic regulation, and DNA damage repair. The most heavily represented complex was SWI/SNF chromatin remodeling, with subunits of both canonical BAF and PBAF enriched.

These included the catalytic ATPases SMARCA4 and SMARCA2; core subunits SMARCC1/2 and SMARCB1; the accessory factors ARID1A and ACTL6A, and the PBAF-specific subunit ARID2. At a common threshold of log2FC > 1 and Benjamini-Hochberg adjusted p < 0.01, 24 proteins were shared between the AR-FL and AR-V7 interactomes (218 and 152 proteins, respectively), half of them published AR interactors and a third SWI/SNF subunits (SI Appendix, Fig. S4B).

Transcriptional elongation and pausing machinery formed a second prominent module, including BRD4 and AFF4, which recruit P-TEFb to gene promoters, together with SUPT5H and NELFA (DSIF and NELF), ELL (a component of the super elongation complex), LARP7, which stabilizes the 7SK snRNP that sequesters P-TEFb in its inactive form (54), the H3K79 methyltransferase DOT1L, and the PAF1 complex subunits PAF1 and CDC73. Additional nuclear associations spanned epigenetic regulation, DNA damage and replication, nuclear receptor coregulation, and nuclear transport. Epigenetic regulators with both activating and repressive functions included the COMPASS components SETD1A and WDR5, the H3K9 methyltransferases EHMT1 and EHMT2, the nuclear corepressors NCOR1 and NCOR2, the NuRD components GATAD2A and MBD3, and the NuA4/TIP60 subunits DMAP1 and EP400. DNA damage and replication factors included MRE11, NBN, RPA2, RPA3, PCNA, TP53BP1 and WRNIP1, consistent with reports that MRE11 and ATR are recruited to ligand-activated AR-FL enhancers during productive transcription (55). Also enriched were the nuclear receptor coactivators NCOA1 and NCOA6 and the nuclear transport components TPR, NUP50 and KPNA3.

By contrast, the AR-V7*^NLSmut^* interactome comprised 158 proteins significantly enriched at the same thresholds, and shifted away from chromatin regulators and toward cytoplasmic networks. The most prominent module was the mRNA deadenylation and decay machinery. Eight subunits of the CCR4-NOT deadenylase complex (CNOT2, CNOT3, CNOT6, CNOT6L, CNOT7, CNOT8, CNOT9 and CNOT10) were significantly enriched, alongside the decapping enzymes DCP1B and DCP2, the decapping activator EDC3, the LSm-domain protein LSM12, and the poly(A)-specific nuclease PAN3. The m6A reader YTHDF1 and the splicing factor SRSF6 were also enriched.

A second module comprised chaperone and protein quality-control factors, including the R2TP/prefoldin-like components RPAP3, PIH1D1 and TTI2, the HSP90 co-chaperone CDC37, the cochaperones DNAJB1 and BAG3, and STUB1/CHIP, an E3 ubiquitin ligase previously implicated in quality control of misfolded or mislocalized AR (56, 57).

Cytoskeletal and mitotic regulators were also prominently represented, spanning four classes: the kinesin motors KIF5B and KIF3B, the augmin/HAUS subunits HAUS2/3/5/8, the γ-tubulin adaptor NEDD1, and the kinetochore protein KNSTRN. SEPTIN9 was also strongly enriched, ranking among the five most significant hits by adjusted p value. Vesicular trafficking proteins were also enriched, including SNX2, SNX5 and SNX6, SEC23A/B, COPE, AP3M1, AP3S1 and ERC1, together with the signaling component IKBKG/NEMO and the Hippo pathway effectors YAP1, AMOTL1 and STK4/MST1. Despite this overall shift, several BAF subunits were also enriched in the AR-V7*^NLSmut^* interactome (SMARCC1/2, SMARCB1, SMARCD2, SMARCD3, SMARCE1, ARID1B, DPF2, BCL7C and SS18), although the catalytic ATPases SMARCA4 and SMARCA2 were not among them.

To directly resolve localization-dependent differences between the two protein environments, we used a linear model with variance-stabilizing normalization that controlled for background and shared interactors (Fig. 4F). Among the top 50 proteins by adjusted p value, annotated nuclear proteins segregated with wild-type AR-V7, whereas cytoplasmic proteins segregated with AR-V7*^NLSmut^*. This direct analysis also resolved factors that had fallen below the fold-change cutoff against the isotype control. The Mediator subunit MED1 was significantly enriched with wild-type AR-V7, whereas the signaling kinase TBK1 and the actin crosslinker FLNA were significantly enriched with AR-V7*^NLSmut^*.

Gene Ontology cellular component enrichment of the 50 most significantly enriched proteins of each interactome, ranked by adjusted p value, further reinforced this separation (Fig. 4G and H). For wild-type AR-V7, enriched terms were nuclear and chromatin-associated. The nuclear protein-containing complex was by far the strongest term, followed by chromosome, chromatin, nucleoplasm, histone deacetylase, SWI/SNF superfamily-type and NuA4 histone acetyltransferase complexes, and the replication fork. In contrast, AR-V7*^NLSmut^* was enriched for cytoplasmic and cytoskeletal compartments: the CCR4-NOT core complex ranked first and P-body second, followed by the HAUS complex, centrosome, microtubule-organizing complex, and microtubule and mitotic spindle structures. Strikingly, no enriched cellular-component term was shared between the wild-type and AR-V7*^NLS^ ^mutant^* rankings. Because each panel used only the 50 most significant proteins of its interactome, several proteins named above lie outside them, including ARID2 in the wild-type ranking and CNOT6, CNOT6L, CNOT7, CNOT8 and HAUS2 in the mutant ranking.

Taken together, these data reveal a striking localization-dependent reorganization of the AR-V7 protein environment. In the nucleus, AR-V7 is associated predominantly with chromatin remodeling, transcriptional, epigenetic and DNA-associated machinery; when retained in the cytoplasm, its protein environment shifts toward mRNA decay, cytoskeletal and trafficking networks, and chaperone quality control systems. Rather than functioning solely as a constitutively active transcription factor, AR-V7 appears to participate in distinct molecular programs on each side of the nuclear envelope.

## Discussion

AR-V7 has been studied primarily as a constitutively active nuclear transcription factor, yet the molecular features that enable its persistent nuclear localization and the consequences of preventing that localization have remained incompletely defined. Here, we identify a multipartite nuclear localization signal spanning the DBD-CE3 junction, uncover a requirement for CE3 in transcriptional activity beyond nuclear accumulation, and show that nuclear and cytoplasmically retained AR-V7 occupy strikingly different protein environments. Together, these findings separate three previously intertwined aspects of AR-V7 biology: nuclear entry, transcriptional competence, and localization-dependent protein associations.

### A variant-specific NLS is created at the DBD-CE3 junction

The defining structural consequence of AR-V7 splicing is the replacement of the AR-FL hinge and ligand-binding domain with the unique CE3 sequence (12, 17). Our data show that this splice junction does more than remove the canonical AR-FL trafficking machinery: it creates a new autonomous nuclear localization signal. The minimal import-competent region spans residues 607-643, and progressive mutagenesis identifies R607, R615, R617, K618 and R642 as the principal basic residues contributing to this multipartite signal (Figs. 1G and 2B, F). Thus, AR-V7 nuclear localization is not encoded by CE3 alone or by the residual DBD alone, but emerges from basic residues distributed across their junction.

This architecture is consistent with the conformational behavior of the 607-643 peptide. AlphaFlow modeling predicts a relatively structured central region surrounding R615/R617/K618 flanked by more flexible termini, particularly toward R642 in CE3 (Fig. 2G, H; SI Appendix, Fig. S3). Rather than defining a single rigid NLS geometry, these models suggest that the AR-V7 signal is conformationally dynamic. Such flexibility could permit spatially separated basic residues to form an import surface in multiple conformations, although direct structural analysis of the NLS in complex with its transport machinery will be required to test this model. Together with our previous demonstration of noncanonical AR-V7 nuclear import (Kim et al., 2022), these findings shift the mechanistic question toward how the DBD-CE3 signal is recognized. Identifying the transport factor or nuclear pore component that engages this multipartite surface is therefore an important next step.

### Nuclear localization and transcriptional competence are separable properties of AR-V7

Our previous work provided an early indication that the AR-V7 DBD has an unexpected role in nuclear localization. D-box mutations (A596T/S597T), which disrupt AR-FL dimerization and transcriptional activity, had distinct effects on AR-FL and AR-V7: in AR-V7, these mutations also caused cytoplasmic sequestration, implicating the DBD in nuclear import (Kim et al., 2022). The present identification of a multipartite NLS spanning the DBD-CE3 junction (Figs. 1G and 2B, F) provides a molecular framework for that observation.

Importantly, the present data also reveal that nuclear localization and transcriptional competence can be uncoupled. Mutants within CE3 retained substantial nuclear localization yet were inactive in the ARE reporter assay (Fig. 2B, C), whereas the strongly import-defective D2/C5 mutant lost the AR-V7 transcriptional program genome-wide (Fig. 3D-F). These data establish nuclear entry as necessary for AR-V7 transcriptional function while raising the possibility that residues within the DBD-CE3 NLS contribute to transcriptional competence once AR-V7 reaches the nucleus.

The mechanism underlying this second function remains unresolved. Mutations in this region could alter DNA binding or chromatin engagement, or CE3 could contribute to the recruitment or stabilization of transcriptional coregulators. The nuclear AR-V7 proximity proteome provides candidates for the latter possibility, including extensive representation of SWI/SNF, NuRD, transcription elongation machinery, and nuclear receptor coregulators (Fig. 4D, G). However, because SWI/SNF components are also recovered in AR-FL interactomes, these associations cannot yet be assigned specifically to CE3. Direct measurements of DNA occupancy and cofactor recruitment by localization-competent CE3 mutants will be required to distinguish these mechanisms. An important caveat is that the current reporter experiments do not independently vary nuclear AR-V7 abundance and NLS sequence. Thus, reduced nuclear concentration could contribute to the loss of reporter activity in partially localized mutants. Nevertheless, the absence of a graded relationship between localization and transcription across the C-series supports investigating a direct role for CE3 in transcriptional competence.

### Cytoplasmic retention exposes a distinct AR-V7 protein network

The cytoplasm has largely been treated as a transit compartment for AR-V7. Compartment-resolved µMap suggests that this view is incomplete. Nuclear AR-V7 was associated predominantly with chromatin remodeling, transcriptional regulation, epigenetic machinery, and DNA-associated proteins, whereas AR-V7^NLSmut^ was enriched for mRNA decay, chaperone, trafficking, and cytoskeletal networks (Fig. 4D-H). Their direct comparison revealed a striking segregation of nuclear and cytoplasmic proteins, demonstrating extensive reorganization of the AR-V7 protein environment when nuclear entry is disrupted (Fig. 4F). AR-V7 has previously been linked to the DNA damage response through changes in repair gene expression and radioresistance (58–60). Our data add a protein-level observation, placing nuclear AR-V7 in proximity to the repair machinery itself. The association with canonical BAF and PBAF SWI/SNF complexes, by contrast, has not been reported for AR-V7. The CCR4-NOT signal is particularly intriguing. Multiple components of this major cytoplasmic deadenylase complex (61) were among the most prominent proteins associated with AR-V7^NLSmut^, and P-body was one of the strongest enriched cellular-component terms (Fig. 4E, H). These observations raise the possibility that cytoplasmically retained AR-V7 may associate with post-transcriptional regulatory machinery. The proximity data do not establish direct interaction or a role for AR-V7 in mRNA decay, as co-residence within P-bodies or related RNA granules could generate the same signature. This possibility is supported by the strong P-body enrichment and a pattern resembling cytoplasmic puncta observed with AR-V7^NLSmut^ by confocal microscopy (Figs. 2E and 4C), although these experiments did not include P-body markers or quantify their co-localization. Notably, a CNOT1 peptide has previously been reported to bind AR-V7 on coregulator arrays (Cato et al., 2019), providing independent evidence for a potential connection between AR-V7 and CCR4-NOT machinery. Distinguishing functional engagement from co-residence will require co-staining with established P-body markers together with measurements of mRNA half-lives or poly(A) tail lengths in the presence of cytoplasmic AR-V7. If this association proves functional, cytoplasmic AR-V7 may influence transcript stability rather than transcription itself, revealing a previously unrecognized post-transcriptional dimension of AR-V7 biology.

Alongside the RNA decay machinery, cytoskeletal and mitotic networks showed strong localization-dependent partitioning (Fig. 4E, F, H). Kinesins and augmin/HAUS components were enriched with AR-V7^NLSmut^, as was SEPTIN9 (Fig. 4E), whereas other cytoskeletal and mitotic proteins, including MAD1L1 and KIF4A, with reported relevance to prostate cancer biology (62 ), preferentially associated with nuclear-competent AR-V7. KIF4A is particularly notable because it has independently been shown to interact with AR-FL and AR-V7 and to stabilize these receptors by inhibiting CHIP-mediated degradation (Cao et al., 2020). KIF4A knockdown restored enzalutamide sensitivity in castration-resistant prostate cancer cells, underscoring its potential contribution to AR-driven resistance. Our data provide independent support for an AR-V7-KIF4A association and suggest that it preferentially associates with nuclear-competent wild-type AR-V7 rather than with the cytoplasm-restricted mutant.

Despite the strong compartmental separation, 14 proteins were nonetheless shared between the AR-V7 and AR-V7^NLSmut^ proximity proteomes, including multiple SWI/SNF subunits (ARID1B, BCL7C, DPF2, SMARCB1, SMARCC1/2, SMARCD2, SMARCE1 and SS18) together with the HSP90 co-chaperone CDC37, the kinetochore protein KNSTRN, the chromatin factor HMG20A and the nucleotide-pool enzyme DUT; the fourteenth shared protein is AR itself, which is the bait (SI Appendix, Fig. S4C). The persistence of SWI/SNF components in both datasets is particularly intriguing given the minimal nuclear accumulation and complete transcriptional inactivity of AR-V7^NLSmut^. One possibility is that AR-V7 encounters SWI/SNF subunits or assembly intermediates in the cytoplasm before nuclear entry, whereas the shared chaperone signal may reflect receptor folding and quality control (C. Liu et al., 2018). Residual nuclear AR-V7^NLSmut^ or effects of the NLS mutations themselves, however, cannot be excluded. Taken together, these data indicate that AR-V7’s protein interaction landscape is shaped primarily by its cellular localization, and that its biological activities may reach well beyond chromatin.

### Implications and Limitations

Two limitations are particularly important for interpreting the proximity data. First, µMap reports spatial proximity rather than direct binding, so the datasets should be viewed as compartment-resolved protein environments rather than collections of validated binary interactors. Second, localization and sequence are coupled in our experimental comparison: AR-V7^NLSmut^ differs from wild-type AR-V7 both in subcellular distribution and in the sequence of its DBD-CE3 NLS. Moreover, AR-V7^NLSmut^ accumulates in the cytoplasm to a degree that wild-type AR-V7 does not normally reach. Some associations may therefore result from the mutations themselves or prolonged cytoplasmic residence rather than representing physiological interactions made by wild-type AR-V7 during transit through the cytosol. The recovery of previously reported AR-associated proteins, including YAP1, supports the biological plausibility of the cytoplasmic dataset but does not establish the functional significance of individual associations. Future experiments that redirect wild-type AR-V7 without altering its NLS sequence will be important for resolving these effects.

### Concluding perspective

Together, our findings identify the DBD-CE3 junction as a functional hub that couples AR-V7-specific nuclear import to transcriptional competence and thereby shapes the cellular protein environment accessible to AR-V7. This creates at least two conceptually distinct opportunities to suppress AR-V7: preventing its nuclear entry or disrupting the CE3-dependent functions required for productive transcription after entry.

Our data strongly support nuclear import as a vulnerability because cytoplasmically retained AR-V7^NLSmut^ loses the AR-V7 transcriptional program (Fig. 3D-F). At the same time, the cytoplasmic proximity proteome cautions that transcriptionally inactive does not necessarily mean biologically inert. If the CCR4-NOT, cytoskeletal, or chaperone associations prove functional, therapies that retain AR-V7 in the cytoplasm redirect its activity rather than simply eliminate it. The compartment-resolved datasets may also inform induced-proximity therapeutic strategies, whose success depends on bringing two proteins together within the same cellular compartment. Regulated induced proximity targeting chimeras (RIPTACs) tether a protein selectively expressed in tumor cells to an essential partner, creating a cytotoxic complex preferentially in target-expressing cells (63), while domain-alteration chimeras (DALTACs) use induced proximity to disrupt productive transcriptional complexes (64). Both require the target and recruited partner to occupy the same compartment. The sharply different nuclear and cytoplasmic AR-V7 protein environments defined here therefore provide a compartment-resolved map of candidate partners for such approaches that cannot be inferred from whole-cell interaction datasets alone (Fig. 4F-H). Whether the DBD-CE3 region itself, or proteins that recognize it, can be pharmacologically engaged remains an important next question.

## Materials and Methods

### Cell Culture

HEK-293 T and LNCaP-C4-2 cell lines were obtained from ATCC. For all AR-V7 localization and ARE reporter experiments, HEK293T cells were cultured in phenol-red and glutamine-free Dulbecco’s Modified Eagle’s Medium (DMEM, Corning 17-205-CV) supplemented with 10% charcoal stripped serum (CSS) (Gibco #12676029) and 1X MycoZap-CL (Lonza #VZA-2012). For lentiviral production, HEK293T cells were cultured in Dulbecco’s Modified Eagle’s Medium (DMEM, Corning 10-013-CV) supplemented with 10% FBS and 1X MycoZap-CL. LNCaP-C4-2 cell lines were cultured in phenol-red and glutamine-free RPMI 1640 (Thermo Scientific #11835055) supplemented with 10% CSS and 1X MycoZap-CL. Mycoplasma detection was tested in all cell lines.

### Plasmid Construction

Doxycycline-inducible expression of EGFP-tagged AR-V7, -AR-V7*^NLSMut^*, or EGFP-tagged AR-FL mutant was generated by subcloning into the lentiviral pCW57 MCS1-P2A-MCS2 (Hygro) tet-on vector using Gateway cloning. pCW57-MCS1-P2A-MCS2 (Hygro) was a gift from Adam Karpf (Addgene plasmid # 80922). Restriction cloning was performed to generate EGFP-tagged AR domains. Briefly, AR domains with SalI and AclI restriction sites were amplified by PCR from a pEGFP-C1-AR-FL plasmid. pEGFP-C1-AR was a gift from Michael Mancini (Addgene plasmid #28235). After purification, PCR products and pEGFP-C1-AR were digested with SalI and AclI, run on a 1% agarose gel (ThermoFisher # 16500500) and backbone/insert fragments were excised from gel and subject to gel purification. T4 ligase was used to ligate AR domains to the pEGFP-C1 backbone (Thermo Scientific # EL0011). 2μL of the ligation reaction was used to transform competent DH5-alpha *E.coli* (Invitrogen #18265017), and colonies were screened by colony PCR, then whole plasmid sequencing (EZ-Plasmid, Genewiz). The construct containing mCherry-tagged GTP-hydrolysis defective Ran mutant, pmCherry-C1-RanQ69L (Addgene plasmid #30309), was a gift from Dr. Jay Brenman (Kazgan et al., 2010) and used for live-cell imaging. pCMV5-AR-V7 and pCMV5-AR-V7 D2/C2 mutant were gifts from Dr. Scott Dehm (Chan et al. 2012). AR-V7 NLS mutant variants were generated using the In-Fusion HD Cloning Kit (Takara Bio) to ensure seamless, directional insertion of designed mutations. The pCMV5-AR-V7 plasmid was digested with KpnI and HindIII to excise the region encoding the C-terminal domain of AR-V7 and linearize the vector. Synthetic double-stranded gene blocks (IDT) containing the desired nuclear localization signal mutations were designed with 15 bp terminal sequences homologous to the vector ends (see Table S1), enabling accurate homologous recombination in the In-Fusion reaction. The In-Fusion reaction mixtures were assembled by combining 5X Enzyme Premix, linearized vector (100 ng), and insert (5 ng), brought to a final volume of 5 µL with sterile Milli-Q water, and incubated at 50 °C for 15 minutes for exonuclease-mediated annealing of homologous ends. The constructs were then transformed into Stellar Competent E. coli (Takara Bio), plated on LB agar supplemented with ampicillin, and incubated overnight at 37 °C. Single bacteria colonies were expanded, and plasmids were purified using MiniPrep columns. Plasmid sequences were fully verified by Sanger sequencing.

### Generation of Stable Cell Lines

We generated the doxycycline-inducible HEK293T and LNCaP-C4-2 (tet-on EGFP-AR-V7, EGFP-ARV7*^NLSMut^*, and AR-FL) by infection with a lentiviral construct (see plasmid constructions section). For lentivirus production, HEK293T cells were seeded in a 10cm dish and transfected with a prepared mix in 1.625 mL DMEM (with no supplements) containing 25 μg of pcw57.1-EGFP-AR-V7-Hygro, -EGFP-AR-V7*^NLSMut^* , or - AR-FL-Hygro. 18.75 μg of psPAX2, 6.25 μg of pMD2.G, and 200 μl of FuGENE HD (Promega). The media was replaced after 4 hours and harvested 48 hours post-transfection. HEK293T and LNCaP-C4-2 cells were plated in individual wells of a six-well plate and transduced with the pcw57.1-EGFP-AR-V7-Hygro, -EGFP-AR-V7*^NLSMut^*, or AR-FL-Hygro lentivirus generated above in complete media containing 5 ug/mL polybrene for 8 hours, then selected with 50-100 ug/mL Hygromycin in complete media.

### Transient Transfections of Plasmid and Drug Treatments

30,000-70,000 cells were plated CellTak-coated on coverslips or CellView® Chambers and incubated at 37C for 24 hours. Cells were transfected with Lipofectamine 3000 within 24 hours of plating. In all live imaging studies, including NTR knockout or RanQ69L, cells were treated with 1.5-3ug/mL of Doxycycline for 4-8 hours or overnight to induce the expression of AR-V7 or AR-FL. To induce the nuclear translocation of AR-FL, cells were co-treated with 1-10nM of R1881

### Antibodies and Reagents

Primary antibodies used include rabbit monoclonal AR-V7 targeting the CE3 domain (Cell Signaling Technology, E308L clone). To detect mutants and for AR-FL µMap pilot assay, Rabbit monoclonal anti-AR antibody targeting the AR N-terminus (Cell Signaling Technology D6F11 Clone) was used. For the AR-V7 µMap, monoclonal anti-GFP from Novus Biologicals (NB600-308) was used. The secondary antibody used to detect AR-V7 or AR-V7 Mutants in immunofluorescence studies is Goat anti Rabbit IgG (H+L) Secondary Antibody, Alexa Fluor 568 (Thermo Fisher Scientific A11011).

### Live Imaging and Immunofluorescence

Cells were imaged before and after fixation with 4% paraformaldehyde (PFA) for 15 minutes at room temperature. Time-points were stored at 4C in PBS after fixation and stained in parallel using the same batch of antibody solution. Samples were permeabilized for 15 minutes in 0.25% Triton-X in PBS, and blocked for 1 hour in 1% BSA, 5% Normal Goat Serum (NGS) at room temperature. Samples were incubated with primary antibody at 1:100-1:250 dilution overnight at 4C. After washing for 10 minutes in PBS, they were incubated with secondary antibody (1:250 dilution) and Hoechst (1:1000) at room temperature for no more than 1-2 hours. After washing for 10-15 minutes in PBS, coverslips were mounted with Fluoromount-GTM aqueous mounting medium and stored at 4C before imaging. Cells stained in suspension for μMap studies were trypsinized 24 hours after transfection with AR vectors (Phenol-red free TryPLE) and pelleted. They were resuspended in fixation buffer (2% PFA in 1X PBS) and inverted continuously at room temperature (RT) for 15 minutes. Cells were pelleted and resuspended in FACS buffer (1X PBS, 2% FBS, 2mM EDTA) 3 times to wash, then resuspended in permeabilization buffer (0.25% Triton-X ,1X PBS) for 15 mins with continuous inversion at RT. After pelleting, cells were inverted in blocking buffer (1X PBS, 2% BSA, 2mM EDTA) for 1 hour at 4C. Cells were inverted in primary antibody solution overnight (1:100). For suspension staining optimization, cells were washed 3X in PBS, pelleted, and resuspended in a solution of 1:250 goat-anti-rabbit Alexa 568 (Invitrogen) for 1 hour at 4C. Cells were washed and added to glass slides with coverslips for confocal imaging.

### Microscopy

Confocal images were acquired using an inverted Nikon Ti2-E CSU-W1 spinning disk microscope (Nikon Instruments) equipped with a CSU-W1 SoRa module (Yokogawa) and an ORCA-Fusion BT sCMOS camera (Hamamatsu). Imaging was performed using either a 40× Plan Apo Lambda air objective (numerical aperture 0.95) or a 60× Plan Apo Lambda oil-immersion objective (numerical aperture 1.42). Z-stack images were collected for each field of view, and extended depth-of-field (EDF) projections were generated using NIS-Elements software (Nikon).

### Image analysis

For nuclear quantification of IF images, the nuclei and cytoplasm of cells expressing GFP-tagged AR were segmented with Cellpose 4.1.1 (65) (66) from the DAPI and phalloidin channels, respectively, and the cytoplasm was defined as the cell mask minus the nuclear mask. Intensities were background-corrected by subtracting the median off-cell intensity of each field, and nuclear localization was quantified as the nuclear fraction (integrated nuclear signal divided by integrated whole-cell signal).

### ARE Reporter Assay

HEK293T cells were transfected with pCMV5-AR-V7 and/or NLS mutants. The gaussia luciferase 6X ARE vector (GS241B-gLuc) was a gift from Dr. Azeem (46). HEK293T cells were infected with lentivirus encoding the gLuc-ARE vector (produced as described above) and subject to antibiotic selection by hygromycin at 50 ug/mL. HEK-gLuc-ARE cells were plated in 96-well plates and transfected with AR-V7 and AR-V7 NLS mutant vectors in quadruplicate. 24 hours post-transfection, the media was collected, and the Secrete-PairTM Gaussia Luciferase detection kit (Pharmacopoeia) was used to detect luciferase signal. The signal was normalized to cell density by reading absorbance of cells at 490nm after adding CellTiter 96® Aqueous solution.

### AlphaFlow Conformational Ensemble Analysis

A 100-conformation ensemble of the AR-V7 NLS region (residues 607 to 643) was generated with the AlphaFlow tool (47) on the Neurosnap web server (https://neurosnap.ai/). Ensembles were analyzed with MDTraj using alpha-carbon (Cα) atoms. All conformations were superposed on the highest-confidence model, and per-residue root-mean-square fluctuation (RMSF) was computed across the aligned ensemble. Pairwise Cα RMSD between all conformations was clustered by hierarchical agglomerative clustering with Ward linkage (SciPy), with the number of clusters chosen by silhouette score across k = 2 to 10, and the distance matrix was embedded in two dimensions by multidimensional scaling (scikit-learn). Cα-Cα and Cζ-Cζ distances between Arg607 and Arg642 were measured for each conformation. One conformer of each cluster was rendered in PyMOL (Schrödinger): AlphaFlow models 8 (Cluster 1), 71 (Cluster 2) and 67 (Cluster 3). Of these, only model 67 is the highest-pLDDT member of its cluster, and the reference model on which all conformations were superposed was not selected. The values printed beneath each structure in Fig. 2G are those of the conformer shown, not cluster means.

### RNA Extraction and Sequencing

Total RNA was isolated from LNCaP-C4-2 cells expressing doxycycline-inducible EGFP-AR-FL, -AR-V7, or AR-V7*^NLSMut^* (±Dox, n=3 replicates per condition) using the RNeasy Mini Kit (Qiagen) according to the manufacturer’s instructions. RNA concentration and purity were assessed by NanoDrop spectrophotometry, and RNA integrity was confirmed using the Agilent 2100 Bioanalyzer prior to submission. High-quality RNA samples were shipped on dry ice to Novogene (Sacramento, CA, USA) for library preparation, sequencing, and bioinformatic processing. Briefly, mRNA was enriched using poly-T oligo-attached magnetic beads, fragmented, and reverse transcribed into cDNA. Libraries were constructed using standard Illumina protocols and sequenced on an Illumina NovaSeq 6000 platform to generate paired-end 150 bp reads. Each sample yielded ∼20-30 million reads. Novogene performed quality control and filtering of raw reads to remove adaptors, low-quality bases, and reads with >5% ambiguous nucleotides. Clean reads were aligned to the human reference genome (GRCh38/hg38) using HISAT2, and transcript abundance was quantified as Fragments Per Kilobase of transcript per Million mapped reads (FPKM). Differential gene expression analysis was conducted using DESeq2, with false discovery rate (FDR) correction applied to generate adjusted p-values (padj). Gene set enrichment analysis (GSEA) was performed on ranked gene lists against curated KEGG and Gene Ontology (GO) databases to identify pathways differentially enriched between AR-V7 and AR-V7 D2/C5 conditions. Separately, gene set variation analysis (GSVA) was performed on the normalized expression matrix against a curated panel of 26 androgen-receptor activity and prostate-lineage gene sets. A per-sample enrichment score was computed for each gene set, and a Δscore was calculated as the score in the doxycycline-induced arm minus the mean score of the matched vehicle-control group. Significance was assessed by one-sample t-test of the three per-replicate Δscores against zero, within each gene set and group. Figure 3F shows four of these gene sets; all 26 are reported in the SI Appendix.

### μMap Assay

Photocatalytic secondary antibodies were prepared by allowing unconjugated goat-anti-rabbit secondary antibody (ThermoFisher, #31212) to react with an iridium photocatalyst via an electrophilic NHS ester, followed by size exclusion chromatography to remove unreacted photocatalyst and N-hydroxy succinimide byproducts following previous literature (67). LNCaP-C4-2 cells expressing doxycycline-inducible EGFP-AR-V7 (n=20) or EGFP-AR-V7*^NLSMut^* (n=20) were grown to 80% confluence in T-150 flasks and AR-V7 expression was induced with 3ug/mL Doxycycline (Enzo Life Sciences #ALX-380-273-G005). After 24 hours, cells were trypsinized, resuspended in 5mL of DMEM complete media, fixed with 2% PFA, permeabilized with 0.25% Triton-X in PBS, blocked with 1% BSA and 5% Normal Goat Serum in PBS, and incubated overnight in low-retention Eppendorf tubes with the primary anti-GFP antibody, n=10 per experiment arm (5μg/mL per replicate, Novus Biologicals NB600-308) or Rabbit IgG isotype control n=10 per experiment arm (5μg/mL per replicate, Jackson ImmunoResearch 011-000-003) overnight at 4C with inversion. Cells were washed with FACS Buffer (2% FBS, 2mM EDTA) 3X at room temperature and incubated with the photocatalytic secondary anti-rabbit antibody in the dark at 4C for 1h. The cells were washed with PBS and incubated with diazirine-PEG3-biotin at 250 μM for 10 minutes while sparging with argon to remove oxygen. Subsequently, samples were irradiated with 440 nm light through a 415nm longpass filter for 20 minutes using a custom-built light setup equipped with a 100W chip-on-board LED. After washing 3X with PBS to remove excess diazirine, cells were resuspended in decrosslinking buffer (600mM Tris pH 8.0, 4% SDS), probe sonicated (5 seconds on, 1 second off, 5 seconds on for 3 cycles), and heated at 95C for 15 minutes. Samples were centrifuged at 20,000G for 10 minutes, and protein supernatant was collected and analyzed by BCA. Samples were reduced and alkylated as described in the literature (67). An equal amount of protein per samples was incubated with streptavidin beads (50 μL Cytiva #30152104010150) overnight, washed 3X sequentially with 1% SDS/PBS, 1M NaCl/PBS, 10% EtOH/PBS, and 50 mM Ammonium Bicarbonate (ABC). After the final wash, beads were pelleted and resuspended in 60μL ABC containing 0.2 ug trypsin and incubated overnight at 37C with continuous inversion. Samples were briefly spun down, and beads were precipitated in a magnetic rack for 5 minutes, liquid transferred to a fresh plate, followed by the addition of 6 μL of 5% formic acid. Samples were centrifuged at 2,500G for 10 minutes and transferred to a new plate to remove precipitated debris. Samples were desalted by Oasis plate (Oasis #186001828BA) following previous literature (67). Samples were resuspended in 40 μL 2% ACN in proteomics H2O +0.1% Formic Acid. Samples were analyzed in data-independent acquisition mode in a timsTOF Pro 2 mass spectrometer. Greater than two peptides enriched in 5 out of 6 samples (strict analysis) were plotted in R relative to the untransfected samples and IgG controls. Significance thresholds: |log2FC| > 1 with Benjamini-Hochberg adjusted p < 1e-4 for Fig. 4D and 4E; -log10 p ≥ 4 with no fold-change cutoff for Fig. 4F; and log2FC > 1 with Benjamini-Hochberg adjusted p < 0.01 for SI Appendix, Fig. S4A-C. For network analysis, interactors passing the Fig. 4D and 4E thresholds in the transfected vs. untransfected DEA file were copied into STRING Database (www.string-db.org) for evaluation of biologically meaningful protein networks and interactor clustering.

## Acknowledgements

This work was supported by the National Cancer Institute, National Institutes of Health (R01CA266704 to P.G.; T32CA062948 to C.C.A. and M.N.; T32CA203702 to U.dC. and M.N.; K99CA312863 to X.C); the National Institute of General Medical Sciences, National Institutes of Health (R35-GM147449 and R35-GM147449-02S1 to J.G.; T32GM141949 to N.E.A.); the U.S. Department of Defense Prostate Cancer Research Program (W81XWH-19-1-0666 to P.G.; PC230548 to X.C.); the Daedalus Fund for Innovation (P.G.); and a Prostate Cancer Foundation Young Investigator Award (25YOUN20 to C.C.A; 25YOUN16 to X.C.).

## Author contributions

U.dC., N.E.A., J.G., and P.G. designed research; U.dC., N.E.A., C.B., M.N., K.K.S. and C.C.A. performed research; J.G. contributed new reagents/analytic tools; U.dC., N.E.A., C.B., X.C., X.K.Z., and J.G. analyzed data; and U.dC., N.E.A., and P.G. wrote the paper. All authors contributed to editing the manuscript and approved the manuscript.

## Classification

Biological Sciences / Medical Sciences.

