## Supplemental Figures for "AR-V7 Utilizes a Noncanonical Nuclear Localization Signal to Sustain Androgen-Independent Nuclear Import and Signaling"

### Figure S1

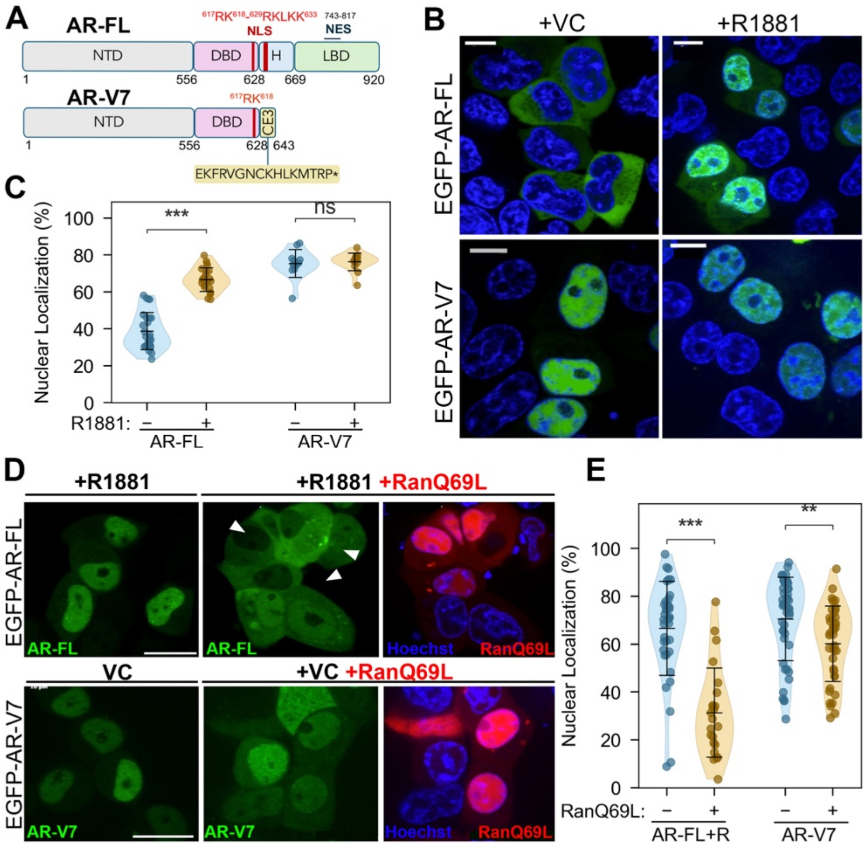


**Figure S1. AR-V7 localizes constitutively to the nucleus largely independently of Ran, in contrast to ligand- and Ran-dependent AR-FL. (A)** Domain architecture of AR-FL and AR-V7. AR-FL contains the NTD, DBD, hinge (H), and ligand-binding domain (LBD), with a bipartite NLS and a nuclear export sequence (NES). AR-V7 retains the NTD and DBD but replaces the hinge, LBD, and NES with the unique 16-residue CE3 sequence (EKFRVGNCKHLKMTRP, ending at residue 643). **(B)** Representative confocal images of HEK293T cells expressing EGFP-AR-FL (top) or EGFP-AR-V7 (bottom) treated with vehicle control (VC) or R1881. Green, EGFP; blue, Hoechst. Scale bars, 10 µm. **(C)** Percentage nuclear localization for EGFP-AR-FL and EGFP-AR-V7 ± R1881. Data points are individual cells overlaid on violin plots; lines indicate mean ± SD. **(D)** Representative confocal images of EGFP-AR-FL (+R1881 alone or +R1881 with dominant-negative mCherry-RanQ69L) and EGFP-AR-V7 (VC alone or VC with mCherry-RanQ69L). Arrowheads indicate cells co-expressing RanQ69L in which AR-FL nuclear accumulation is strongly reduced. Right panels show the merge of Hoechst (blue) and RanQ69L (red). Scale bars, 10 µm. **(E)** Percentage nuclear localization with and without mCherry-RanQ69L as in (C). RanQ69L reduced AR-FL nuclear accumulation from 66% to 31%, while reducing AR-V7 only from 70% to 60%. For (C) and (E), significance was assessed by two-sided Mann-Whitney U test with Bonferroni correction for two comparisons within each panel; n indicates individual cells (C: 26, 24, 12 and 14 cells for AR-FL -R, AR-FL +R, AR-V7 -R and AR-V7 +R, respectively;

E: 38, 23, 45 and 40 cells for AR-FL +R, AR-FL +R with RanQ69L, AR-V7 and AR-V7 with RanQ69L, respectively). *$p < 0.05$; **$p < 0.01$; ***$p < 0.001$; ****$p < 0.0001$; ns, not significant.

**Figure S2**

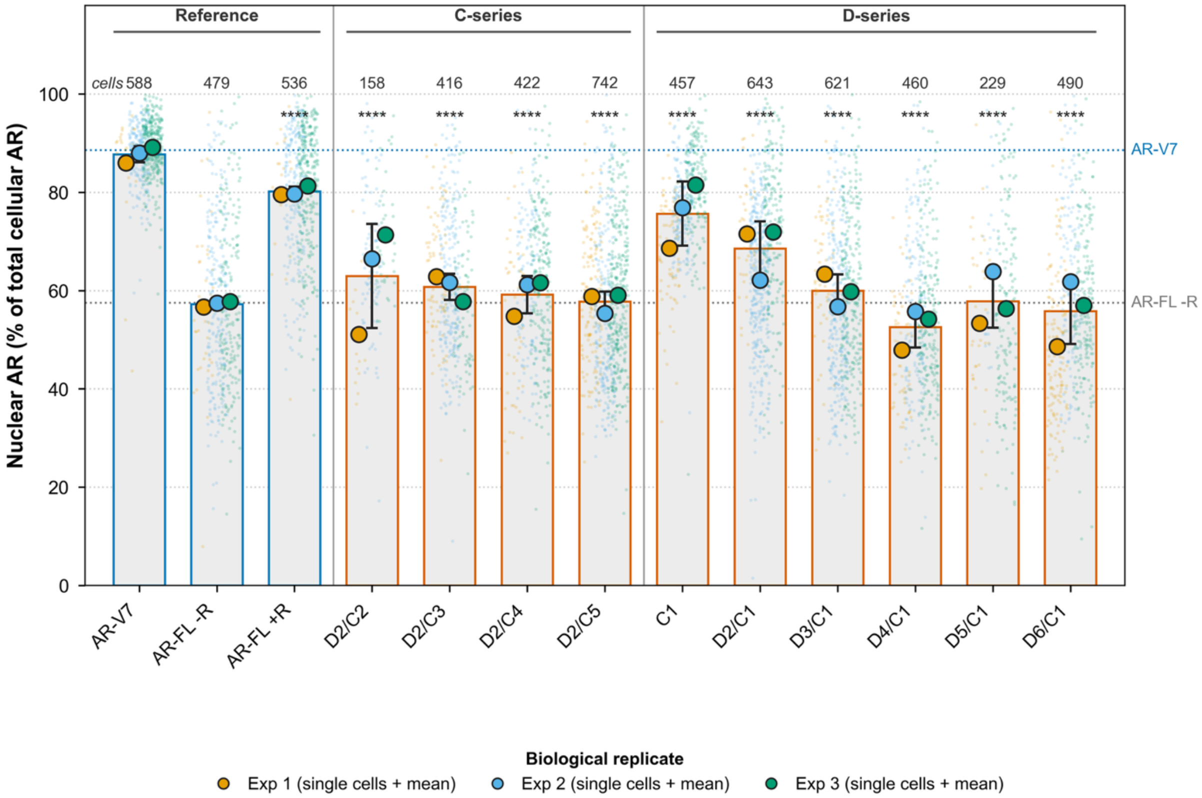


**Figure S2. Raw single-cell nuclear AR across the NLS alanine-mutant panel.** SuperPlot of nuclear AR in HEK293T cells expressing the indicated EGFP-tagged AR constructs, measured by quantitative confocal microscopy. Nuclear AR is the background-corrected integrated EGFP-AR signal within the DAPI-defined nucleus as a percentage of the whole-cell signal, shown here without the normalization applied in Fig. 2B. Small dots, individual cells colored by experiment; large dots, the mean of each of 3 independent experiments; bars, mean of those 3 means ± SD. Cell counts are printed above each column. Significance versus AR-V7 was determined using simultaneous tests of general linear hypotheses (Bonferroni-adjusted) applied to a linear mixed-effects model with genotype as a fixed effect, experiment as a random intercept, and genotype-specific variances, as in Fig. 2B; every construct differed from AR-V7 at ****$p < 0.0001$.

**Figure S3**

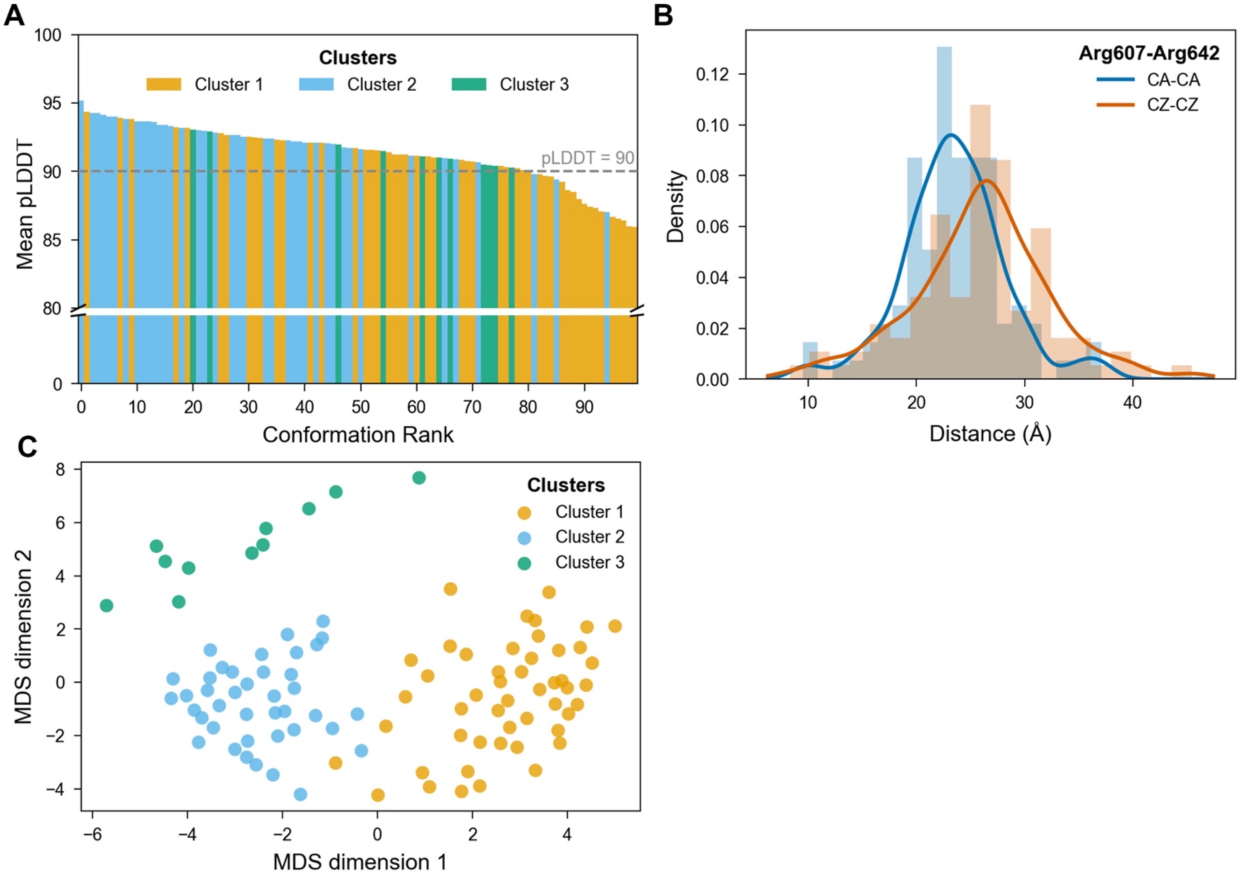


**Figure S3. Conformational ensemble analysis of the AR-V7 NLS peptide. (A)** Per-conformation mean pLDDT across the 100-model AlphaFlow ensemble of the AR-V7 NLS peptide (aa 607-643), ranked from highest to lowest confidence. Bars are colored by conformational cluster (Cluster 1, orange, n = 49; Cluster 2, blue, n = 40; Cluster 3, green, n = 11). The dashed line marks pLDDT = 90; 81 of 100 models exceeded this threshold. **(B)** Density distribution of the Arg607-Arg642 end-to-end distance across the ensemble, measured between Cα atoms (blue) and between terminal guanidinium Cζ atoms (orange), illustrating a broad continuum of compact-to-extended states (Cα-Cα 23.48 ± 4.76 Å; Cζ-Cζ 25.97 ± 6.32 Å; mean ± SD across the 100 conformations). (C) Multidimensional scaling (MDS) projection of pairwise structural distances among the 100 conformations, with each point colored by cluster (colors and cluster sizes as in A), showing three distinct conformational populations.

### Figure S4

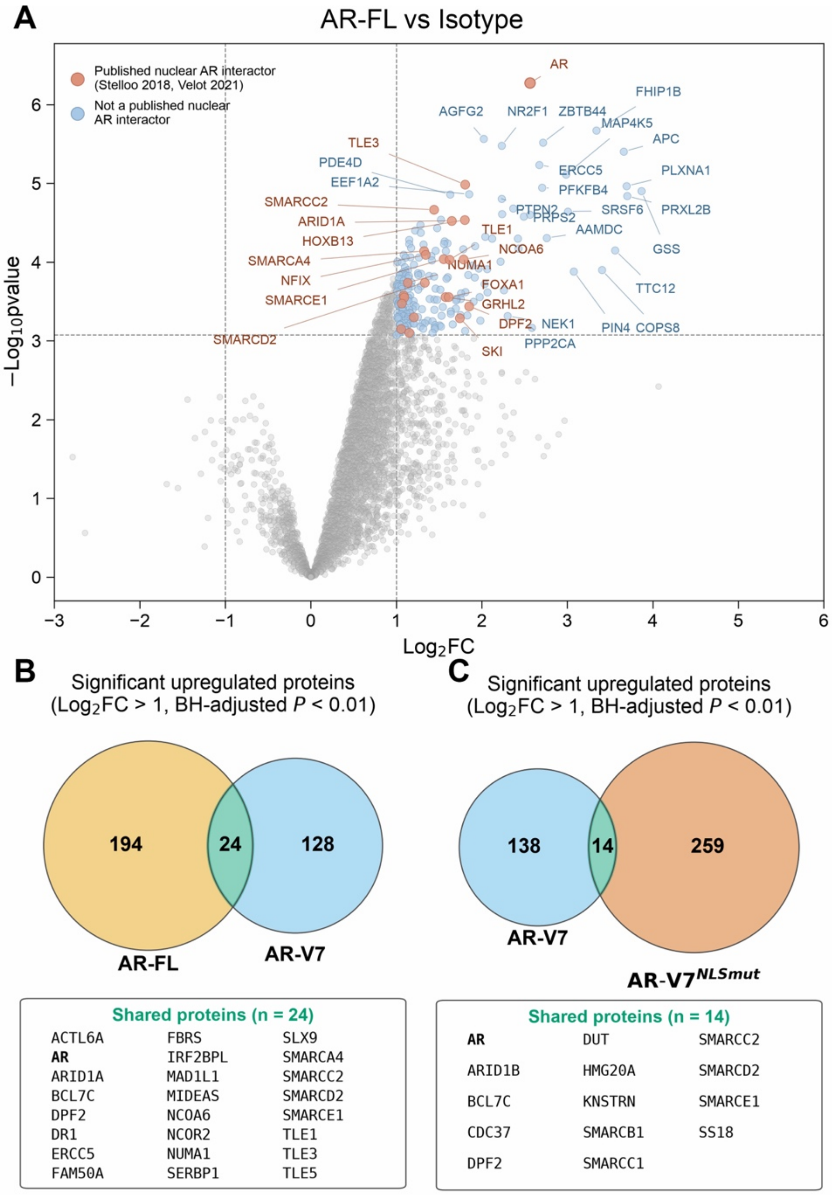


**Figure S4. µMap validation against the AR-FL interactome and shared/distinct interactors across AR isoforms. (A)** Volcano plot of µMap-enriched proteins for AR-FL versus isotype control. Dashed lines indicate the significance thresholds, log2FC > 1 and Benjamini-Hochberg adjusted p < 0.01 (1% false discovery rate); the horizontal line is drawn at the equivalent nominal p value. Salmon, proteins previously reported as nuclear AR interactors; light blue, proteins not previously reported as nuclear AR interactors; grey, non-significant. AR was the most significantly enriched protein, validating the assay. **(B)** Venn diagram of proteins significantly enriched at log2FC > 1 and Benjamini-Hochberg adjusted p < 0.01, shared between the AR-FL and AR-V7 µMap experiments. At this threshold the AR-V7 set contains 152 proteins and the AR-V7$^{NLSmut}$ set 273, compared with 129 and 158 at the stricter adjusted p < 1e-4 cutoff used in Fig. 4D and 4E.

**(C)** Venn diagram of significantly enriched proteins (thresholds as in B) shared between the AR-V7 and AR-V7$^{NLSmut}$ experiments.

**Movie legend:**

**Movie S1. Conformational dynamics of the AR-V7 nuclear localization signal.** Animation of the AlphaFlow conformational ensemble of AR-V7 residues 607 to 643, rendered in PyMOL. The peptide backbone is shown as a blue cartoon beneath a translucent molecular surface. Basic residues of the structured central core are shown as orange sticks, and the Arg607 and Arg642 termini are highlighted in yellow. The central core around residues 614 to 619 stays comparatively fixed, while the termini sample a wide range of positions, consistent with the per-residue fluctuations in Fig. 2H.

### Table S1. Gene Block Primer Sequences for AR-V7 NLS Mutants

| Name | Mutations | Gene block sequence |
|---|---|---|
| D2/C3 | R617A/K618A/K629A/R631A/K 636A | tgtggagatgaagcttctgggtgtcactatggagctctcacatgtggaagctgcaaggtcttcttc aaaagagccgctgaagggaaacagaagtacctgtgcgccagcagaaatgattgcactattg ataaattccgaaggaaaaattgtccatcttgtcgtcttgcggcatgttatgaagcagggatgactc taggagaagcattcgcggttggcaattgcgcgcatctcaaaatgaccagaccctgatctagag gatcccgggtggcat |
| D2/C4 | R617A/K618A/K629A/R631A/K 636A/K639A | tgtggagatgaagcttctgggtgtcactatggagctctcacatgtggaagctgcaaggtcttcttc aaaagagccgctgaagggaaacagaagtacctgtgcgccagcagaaatgattgcactattg ataaattccgaaggaaaaattgtccatcttgtcgtcttgcggcatgttatgaagcagggatgactc taggagaagcattcgcggttggcaattgcgcgcatctcgcgatgaccagaccctgatctagag gatcccgggtggcat |
| D2/C5 | R617A/K618A/K629A/R631A/K 636A/K639A/R642A | tgtggagatgaagcttctgggtgtcactatggagctctcacatgtggaagctgcaaggtcttcttc aaaagagccgctgaagggaaacagaagtacctgtgcgccagcagaaatgattgcactattg ataaattccgaaggaaaaattgtccatcttgtcgtcttgcggcatgttatgaagcagggatgactc taggagaagcattcgcggttggcaattgcgcgcatctcgcgatgaccgcaccctgatctagag gatcccgggtggcat |
| D0/C1 | R642A | tgtggagatgaagcttctgggtgtcactatggagctctcacatgtggaagctgcaaggtcttcttc aaaagagccgctgaagggaaacagaagtacctgtgcgccagcagaaatgattgcactattg ataaattccgaaggaaaaattgtccatcttgtcgtcttcggaaatgttatgaagcagggatgact ctaggagaaaaattccgggttggcaattgcaagcatctcaaaatgaccgcaccctgatctaga ggatcccgggtggcat |
| D2/C1 | R617A/K618A/ R642A | tgtggagatgaagcttctgggtgtcactatggagctctcacatgtggaagctgcaaggtcttcttc aaaagagccgctgaagggaaacagaagtacctgtgcgccagcagaaatgattgcactattg ataaattccgaaggaaaaattgtccatcttgtcgtcttgcggcatgttatgaagcagggatgactc taggagaaaaattccgggttggcaattgcaagcatctcaaaatgaccgcaccctgatctagag gatcccgggtggcat |
| D3/C1 | R615A/ R617A/K618A/ R642A | tgtggagatgaagcttctgggtgtcactatggagctctcacatgtggaagctgcaaggtcttcttc aaaagagccgctgaagggaaacagaagtacctgtgcgccagcagaaatgattgcactattg ataaattccgaaggaaaaattgtccatcttgtgcgcttgcggcatgttatgaagcagggatgact ctaggagaaaaattccgggttggcaattgcaagcatctcaaaatgaccgcaccctgatctaga ggatcccgggtggcat |
| D4/C1 | R607A/ R615A/ R617A/K618A/ R642A | tgtggagatgaagcttctgggtgtcactatggagctctcacatgtggaagctgcaaggtcttcttc aaaagagccgctgaagggaaacagaagtacctgtgcgccagcagaaatgattgcactattg ataaattcgcaaggaaaaattgtccatcttgtgcgcttgcggcatgttatgaagcagggatgact ctaggagaaaaattccgggttggcaattgcaagcatctcaaaatgaccgcaccctgatctaga ggatcccgggtggcat |
| D5/C1 | R607A/R608A/R615A/ R617A/K618A/ R642A | tgtggagatgaagcttctgggtgtcactatggagctctcacatgtggaagctgcaaggtcttcttc aaaagagccgctgaagggaaacagaagtacctgtgcgccagcagaaatgattgcactattg ataaattcgcagcaaaaaattgtccatcttgtgcgcttgcggcatgttatgaagcagggatgact ctaggagaaaaattccgggttggcaattgcaagcatctcaaaatgaccgcaccctgatctaga ggatcccgggtggcat |
| D6/C1 | R607A/ R608A/K609A/R615A/ R617A/K618A/ R642A | tgtggagatgaagcttctgggtgtcactatggagctctcacatgtggaagctgcaaggtcttcttc aaaagagccgctgaagggaaacagaagtacctgtgcgccagcagaaatgattgcactattg ataaattcgcagcagcaaattgtccatcttgtgcgcttgcggcatgttatgaagcagggatgact ctaggagaaaaattccgggttggcaattgcaagcatctcaaaatgaccgcaccctgatctaga ggatcccgggtggcat |
